# Syndri: Context-aware removal of transcript spillover in imaging-based spatial transcriptomics

**DOI:** 10.64898/2026.11.24.751906

**Authors:** Jeff Winchell, Jian Ma, Spencer Krieger

## Abstract

Imaging-based spatial transcriptomics (ST) measures gene expression at single-cell resolution, but imperfect segmentation often misassigns transcripts between neighboring cells, a problem known as transcript spillover. This spillover is niche-dependent and can therefore confound downstream analyses that rely on niche-specific differential expression. True spillover cannot be directly quantified in experimental ST data, so existing correction methods infer contamination indirectly from transcript positions or expression structure. Here we introduce Syndri, a post-segmentation correction method that frames spillover detection as a supervised, gene-level classification task. We develop Syndri-Sim to generate labeled training data by simulating spillover between spatially mapped scRNA-seq profiles. Syndri uses a transformer to predict gene-level spillover probabilities conditioned on each cell’s expression, identity, and local neighborhood, without requiring transcript-level coordinates or deconvolution. On held-out simulated data from Syndri-Sim, Syndri achieves an AUPRC of 0.86, compared with 0.64 for the next-best method. Across MERFISH, CosMx, and Xenium datasets, Syndri achieves the highest aggregate performance across metrics of cell-type separation, reference similarity, and preservation of within-cell-type heterogeneity. In a CosMx lung cancer dataset, Syndri removes nearly all potentially contaminating tumor and macrophage markers from differential expression between tumor-associated and stromal fibroblasts while retaining native fibroblast markers. These results demonstrate the utility of learning from simulated spillover to improve the interpretation of spatial gene expression.

## Introduction

Imaging-based spatial transcriptomics (ST) measures the locations of individual RNA transcripts within intact tissue sections, enabling single-cell quantification of gene expression while preserving spatial context [1–3]. Platforms such as Xenium [3], CosMx [2], and MERFISH/MERSCOPE [1] are now widely used to study tissue organization and how local microenvironments shape cellular identity and function [4]. Many of these studies rely on differential expression (DE) analysis to identify spatial transcriptional programs and characterize interactions between neighboring cell populations [5–8]. The reliability of these conclusions therefore depends on the accuracy of cell-level transcript counts.

However, cell-level counts are often confounded by transcript spillover, in which transcripts from one cell are assigned to a neighboring cell due to segmentation errors, optical crowding, or transcript diffusion [9, 10]. Spillover distorts cell-type-specific expression profiles, introduces spurious DE genes [11], and can obscure true differences between cell populations [9, 12]. Because contaminating transcripts come from adjacent cells, the spillover a cell receives depends on its microenvironment. As a result, spillover is particularly problematic when comparing cells of the same type across different niches, which is central to many spatial analyses [7, 13]. For example, nonmalignant cells adjacent to tumor cells may appear to express tumor-associated markers due to transcript misassignment instead of endogenous expression [12, 14].

Several computational methods have recently been developed to correct spillover after segmentation. Because ground-truth spillover cannot be measured directly in ST data, these methods infer contamination without supervision, relying on assumptions about how spillover appears in the data. cellAdmix [12] and ovrlpy [15] use transcript coordinates to identify spillover, assuming that misassigned transcripts are spatially co-located. These approaches require transcript-level coordinates and rely on the spatial distribution of transcripts to identify contamination. However, contaminating transcripts can be distributed throughout recipient cells rather than being confined to their boundaries (**Fig**. 1b), potentially complicating their distinction from endogenous expression. SPLIT [9] uses the deconvolution method RCTD [16] to estimate cell-type mixtures and then rescales counts based on a scRNA-seq reference, so its corrections depend on deconvolution accuracy and do not directly model each cell’s spatial neighborhood. ResolVI [17] uses a latent-variable model to separate intrinsic expression from contamination and reconstructs corrected expression from its posterior instead of directly modifying observed counts. In each case, correction accuracy depends on how well the method’s assumptions hold. There remains a need for a method that learns spillover patterns from labeled examples, accounts for each cell’s local neighborhood, and removes contamination while preserving biological variation within cell types.

**Figure 1:**
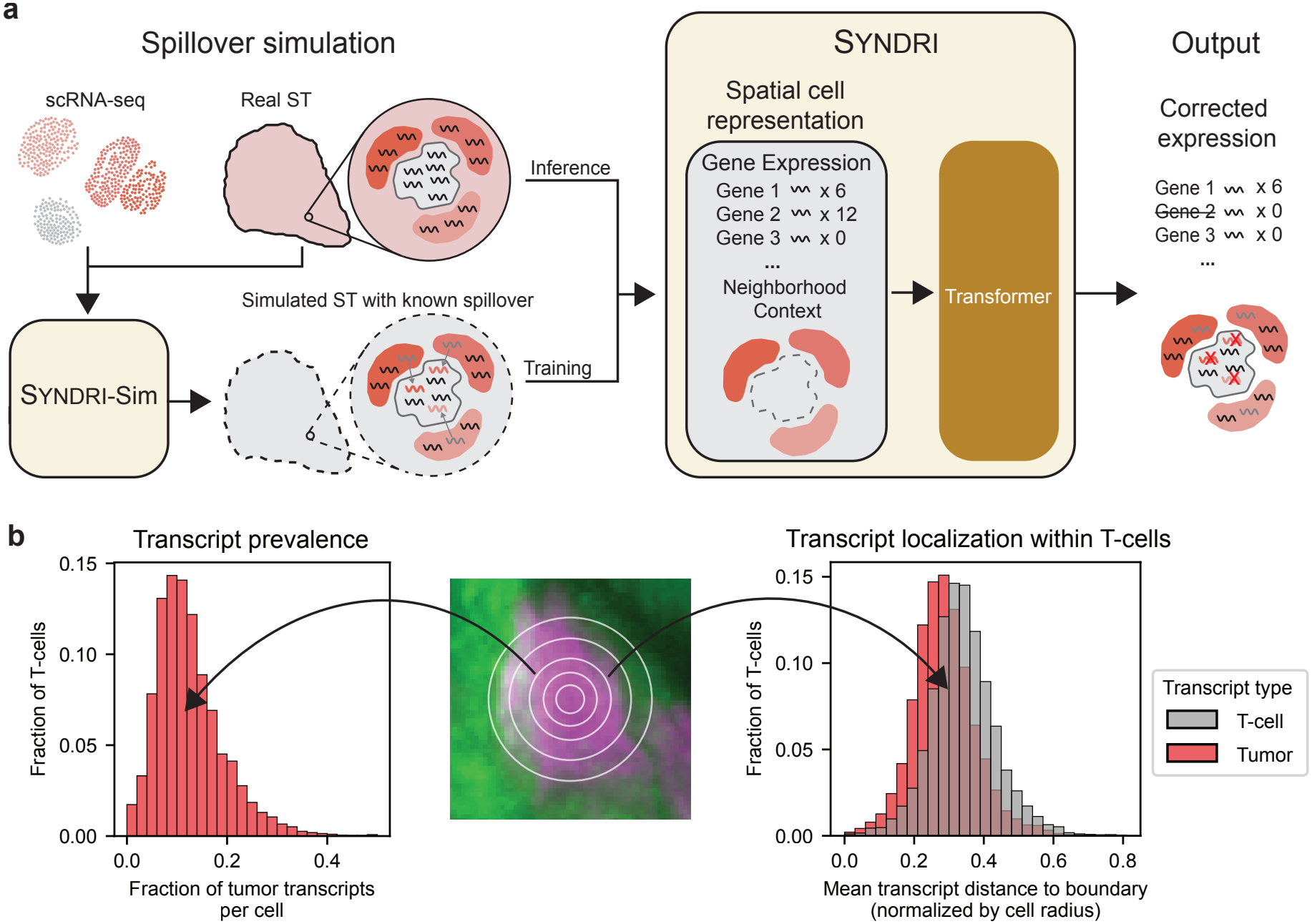
Overview of Syndri. **a**. Syndri frames spillover correction as supervised learning. During training, Syndri-Sim assigns spillover-free scRNA-seq profiles to the cell positions of a real ST sample and transfers transcripts between neighboring cells, producing simulated ST data with known spillover. Syndri takes each cell’s gene expression and the identities of its neighboring cells and predicts which genes contain spillover. At inference, the trained model is applied to real ST data, and transcripts of genes predicted to be contaminated are removed from the observed counts. **b**. Histograms showing the distribution of the fraction of each T cell’s transcripts that come from tumor-marker genes (left) and the distribution of mean distance of transcripts to the cell boundary (right). Tumor transcripts lie closer to the boundary than T cell transcripts, but are often found near the cell center.

Here we present Syndri, a method that corrects transcript spillover by reframing it as a supervised learning problem. Syndri generates labeled training data by assigning spillover-free scRNA-seq profiles to the cell positions of an ST sample, producing a synthetic tissue with known expression. It then simulates spillover by transferring transcripts between neighboring cells in this synthetic tissue. A transformer trained on these data uses each cell’s neighborhood to identify spillover specific to its local context. Syndri operates directly on cell-by-gene count matrices, does not require transcript coordinates or cell-type deconvolution, and only removes transcripts from the observed counts instead of resampling them. Across datasets spanning various species, tissues, disease contexts, and ST platforms we show that Syndri outperforms existing spillover correction methods in cell-type separation, similarity to reference expression profiles, and preservation of within-cell-type heterogeneity. We also show Syndri outperforms other methods directly using our simulated data with known spillover as a benchmark. In a CosMx non-small cell lung cancer dataset, Syndri was the only method that removed spillover-driven tumor and macrophage markers from the genes differentially expressed between fibroblasts in tumor and stromal regions.

## Results

### Overview of Syndri

Syndri corrects transcript spillover by learning from examples in which the spillover is known. Although spillover arises through several mechanisms, each adds transcripts from neighboring cells to a cell’s measured expression profile (**Fig**. 1b). Endogenous and spillover transcripts contribute to the same observed counts and cannot be distinguished from those counts alone, so existing methods rely on fixed assumptions about how spillover appears. In contrast, scRNA-seq profiles are free of spatial spillover because cells are dissociated before sequencing, although they can contain ambient RNA [18, 19]. Spillover can therefore be simulated in these profiles with known ground truth. This moves assumptions about spillover from the model to the simulation, where they can be inspected and varied, and lets the model learn spillover patterns that are not explicitly specified. Simulation also provides an effectively unlimited supply of training examples for high-capacity architectures such as transformers.

To generate synthetic ST data with known spillover, Syndri-Sim assigns scRNA-seq profiles to the cell positions of the ST dataset by matching cell types, producing a synthetic tissue (**Fig**. 1a). It then adds transcripts from each query cell’s spatial neighbors to its expression profile, so each simulated cell has known gene-level labels indicating which genes received spillover. We frame correction as gene-level binary classification: a transformer is trained on these examples to predict the probability that each expressed gene contains spillover, given the cell’s expression profile, its cell type, and the cell types of its neighbors. The trained model is then applied to the real ST data. Within each annotated cell type and spatial slice, cell–gene entries are removed in descending probability order until a target fraction of transcripts has been removed, subject to a minimum probability threshold.

### Syndri improves cell-type separation while preserving reference similarity and within-cell-type heterogeneity

We benchmarked Syndri against SPLIT [9], ResolVI [17], ovrlpy [15], and cellAdmix [12], along with the uncorrected data, on whole mouse brain MERFISH (BM), non-small cell lung cancer CosMx (NC), and breast cancer Xenium (BX) datasets (**Tab**. 1). ovrlpy and cellAdmix require transcript-level coordi-nates and could therefore be evaluated only on NC and BX (gray entries in **Fig**. 2b). Because spillover correction can fail in distinct ways, we scored each method on three criteria (**Fig**. 2): cell-type separation, which should improve as cross-cell-type contamination is removed; reference similarity to a matched, comparatively uncontaminated single-cell reference; and preservation of within-cell-type heterogeneity, which guards against over-smoothing. We ranked methods on each metric per slice, averaged ranks across slices within each dataset, and weighted the three criteria equally in the cross-metric relative-rank score.

**Table 1.** Summary of spatial transcriptomics datasets and matched single-cell references used in this study.

| Abbrev. | Dataset | Technology | Spatial cells | Genes | Reference cells | Spatial sections |
|---|---|---|---|---|---|---|
| BM | Whole mouse brain | MERFISH | 215,278 | 1,122 | 271,164 | 3 |
| NC | NSCLC | CosMx | 771,236 | 956 | 40,362 | 8 |
| BX | Breast cancer | Xenium | 1,673,926 | 277 | 10,658 | 19 |

**Figure 2:**
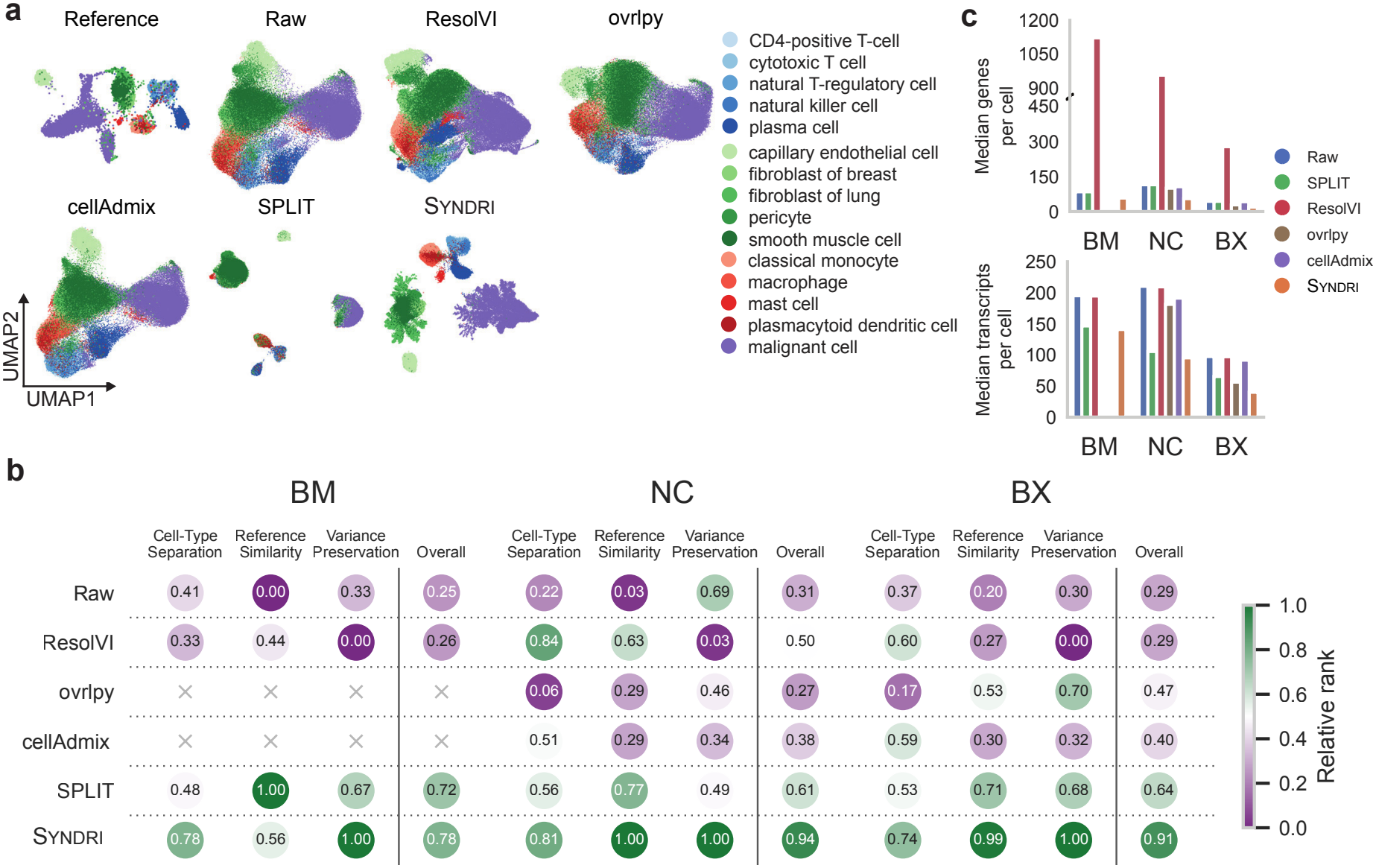
Benchmarking of spillover correction methods. **a**. UMAP [20] embeddings of the breast cancer Xenium (BX) scRNA-seq reference and of a single spatial slice, uncorrected (Raw) and after correction by each method, colored by cell type. **b**. Relative rank of correction methods for three datasets under three criteria: cell-type separation, cosine similarity to the matched single-cell reference (Reference Similarity), and Spearman correlation of gene-wise variance with the reference (Variance Preservation). Ranks were computed per slice and averaged across slices within each dataset, and Overall is the unweighted mean of the three criteria. Gray crosses mark evaluations that could not be performed. **c**. Median number of detected genes (top) and transcripts (bottom) per cell for each method in each dataset.

Relative to the uncorrected data, both SPLIT and Syndri visibly improved cell-type separation in BX (**Fig**. 2a). However, SPLIT retained overlap between several cell populations, including fibroblast-associated structure that was discordant with the matched single-cell reference annotations. This overlap likely reflects SPLIT’s reliance on RCTD deconvolution: where RCTD assigns ambiguous or incorrect cell-type mixtures to a cell, SPLIT’s rescaling inherits those errors (**Fig**. S1). Syndri, which does not depend on deconvolution, produced clearer separation while maintaining overall correspondence with the matched single-cell reference. In contrast, ResolVI, ovrlpy, and cellAdmix showed little change in cell-type separation relative to the uncorrected data.

Quantitatively, Syndri achieved the highest aggregate score in all three datasets, with the largest margins over competing methods in the two cancer datasets, NC and BX (rankings in **Fig**. 2b; metrics in **Fig**. S2). It ranked first or second for cell-type separation in every dataset and first for heterogeneity preservation in every dataset. All correction methods improved reference similarity relative to the uncorrected data, consistent with spillover shifting spatial expression profiles away from the matched single-cell reference. Syndri achieved the highest reference similarity in NC and BX, whereas SPLIT ranked first in BM.

ResolVI displayed a different performance profile. Although it frequently achieved high silhouette scores, it performed substantially worse on ARI and NMI [21] and consistently showed the lowest heterogeneity preservation (**Fig**. S2c). This pattern follows from its generative design: ResolVI reconstructs expression by sampling from a learned posterior, so many originally zero entries receive small fractional values, which increases the number of detected genes per cell while leaving total transcript abundance similar (**Fig**. 2c) [17]. After library-size normalization and log transformation, these small values can contribute appreciably to downstream representations, inflating silhouette scores while degrading within-cell-type structure [22]. This discrepancy illustrates why cell-type separation alone is insufficient to evaluate correction quality: a method can produce compact, well-separated embeddings while substantially altering within-cell-type expression structure.

Overall, competing methods each traded one criterion against another: ResolVI sacrificed within-cell-type structure for separation, and ovrlpy and cellAdmix yielded limited gains in separation. SPLIT’s performance was potentially sensitive to differences in cell-type assignments obtained through deconvolution. By learning spillover patterns from labeled synthetic examples and only removing counts, Syndri was the only method to perform consistently well on all three criteria across species, tissues, and platforms.

### Syndri outperforms existing methods on simulated data with known spillover

Because ground-truth spillover cannot be measured in ST data, the benchmarks above rely on indirect measures. Syndri-Sim builds simulated ST data from spillover-free scRNA-seq profiles and therefore records the true spillover of every gene in every cell, so it can also serve as a ground-truth benchmark. We first asked how closely the simulated data resemble real ST data in the NC dataset. Adding spillover with Syndri-Sim reduced the RBF MMD^2^ [23] between cell-level expression distributions within each cell type of the single-cell data and the matched spatial counts used to define the cell layout by around 30% across all three replicates (**Fig**. 3a). By retaining the cell positions and annotations of the original spatial section, Syndri-Sim preserves the observed cell-type adjacency structure by construction (**Fig**. 3b).

**Figure 3:**
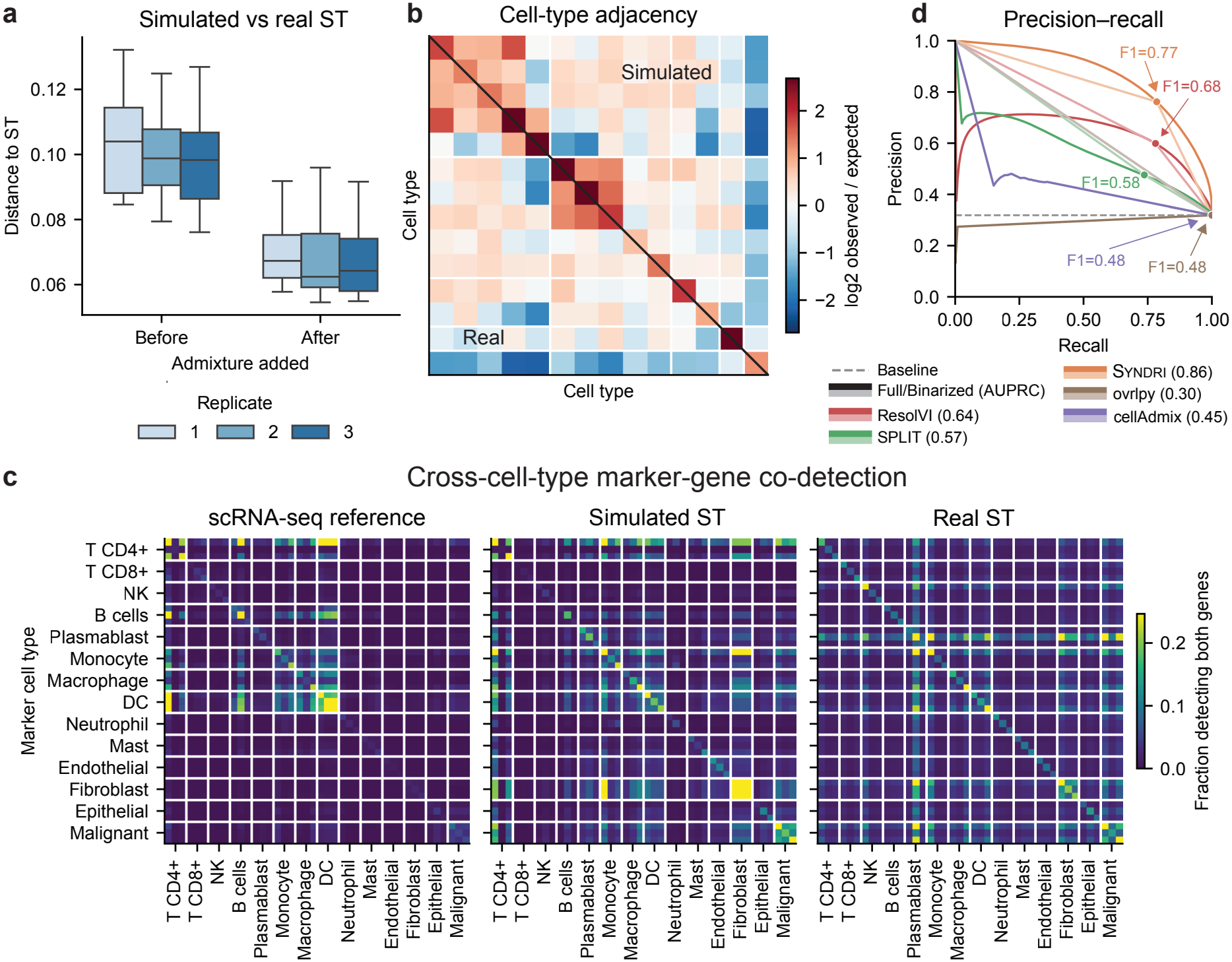
Evaluation of spillover correction on simulated spatial data with known ground truth from Syndri-Sim. **a**. Distance (RBF MMD^2^) between mapped scRNA-seq profiles or simulated ST profiles and matched real ST expression, with the replicates shown evaluated for simulator parameter selection. **b**. Cell-type adjacency in a single slice of real ST data (lower triangle) and simulated ST data (upper triangle), shown as the log_2_ ratio of the observed frequency of each neighbor cell type among the neighbors of each query cell type to the frequency expected from overall cell-type abundance. **c**. Co-detection frequencies for pairs of marker genes, using the top three markers per cell type. Genes are grouped by their associated cell type along both axes, and each heatmap entry represents the fraction of cells in which both genes were detected. **d**. Precision–recall curves for Syndri, SPLIT, ResolVI, cellAdmix, and ovrlpy at identifying simulated spillover across four held-out test slices. Dark curves show continuous contamination scores, with AUPRC values in the legend. For Syndri, these scores are predicted admixture probabilities; for comparison methods, they are removal scores calculated as 1 − *c/a*, where *c* and *a* denote corrected and admixed counts. Light curves show binary predictions using oracle thresholds selected to maximize F1-score against known contamination labels, with the corresponding F1-scores annotated. The dashed line indicates the prevalence-based random baseline.

This allows contamination to be simulated within realistic tissue neighborhoods, although adjacency agreement does not independently validate the contamination model. We also compared the co-detection of marker genes from different cell types, which should be rare in uncontaminated cells (**Fig**. 3c). In the scRNA-seq reference, such co-detection was largely limited to related immune populations, whereas the real ST slice showed much broader co-detection, particularly involving fibroblast, epithelial, and malignant markers. Syndri-Sim introduced co-detection in these same groups of markers, though it did not reproduce every pattern in the ST slice. This gap may arise because real spillover also comes from cells above and below the imaged plane [15], whereas Syndri-Sim transfers transcripts only between neighboring cells in two dimensions, so the simulated and real data can match in adjacency yet differ in co-detection.

We used Syndri-Sim to compare Syndri with SPLIT, ResolVI, ovrlpy, and cellAdmix. To evaluate methods requiring transcript-level coordinates, we generated synthetic transcript positions within the observed segmentation masks, retaining the known origin of each transcript. The NC single-cell reference contains data from seven donors. We simulated training data from five donors for Syndri, used one for validation, and evaluated all methods on simulated data from the remaining donor. SPLIT used the combined training and validation donors as its reference, while ResolVI was fitted independently to each test slice.

Syndri-Sim adds spillover only to genes that the query cell does not endogenously express. We therefore evaluated contamination detection over expressed cell–gene entries using continuous removal scores and binarized predictions. For Syndri, continuous scores were its predicted admixture probabilities; for methods returning corrected counts, scores were calculated as *s* = 1 *− c/a*, where *c* and *a* denote corrected and admixed counts. Continuous scores were evaluated using AUPRC, while binary predictions were evaluated using F1-score at oracle thresholds chosen to maximize F1 against the simulated ground truth (**Fig**. 3d).

Syndri achieved the highest AUPRC (0.86), compared with ResolVI (0.64), SPLIT (0.57), cellAd-mix (0.45), and ovrlpy (0.30). At their respective oracle operating points, Syndri also achieved the highest F1-score (0.77), compared with ResolVI (0.68), SPLIT (0.58), and ovrlpy and cellAdmix (both 0.48) (**Fig**. 3d).

This benchmark has limitations. Syndri is trained on data from the same simulator, although on different donors, so it is evaluated on spillover of the type it was trained to remove. Because spillover is added only to genes not endogenously expressed by the query cell, the task favors binary detection, which does not match how ResolVI and SPLIT adjust counts; we therefore report both real-valued and binarized scores. Additionally, the synthetic transcript-coordinate model approximates subcellular localization using uniform native-transcript placement and donor-biased contamination, which may not capture the spatial patterns exploited by transcript-level correction methods. Syndri-Sim also does not fully reproduce the marker co-expression of real ST data. Nevertheless, the simulation captures key co-expression patterns of real ST data and gives a setting with known spillover, in which Syndri substantially outperformed existing methods.

### Syndri enables more reliable downstream biological analyses

Spatial differential expression (DE) analysis underlies many downstream analyses, so we tested whether spillover correction improves its conclusions in the NC dataset (**Fig**. 4).

**Figure 4:**
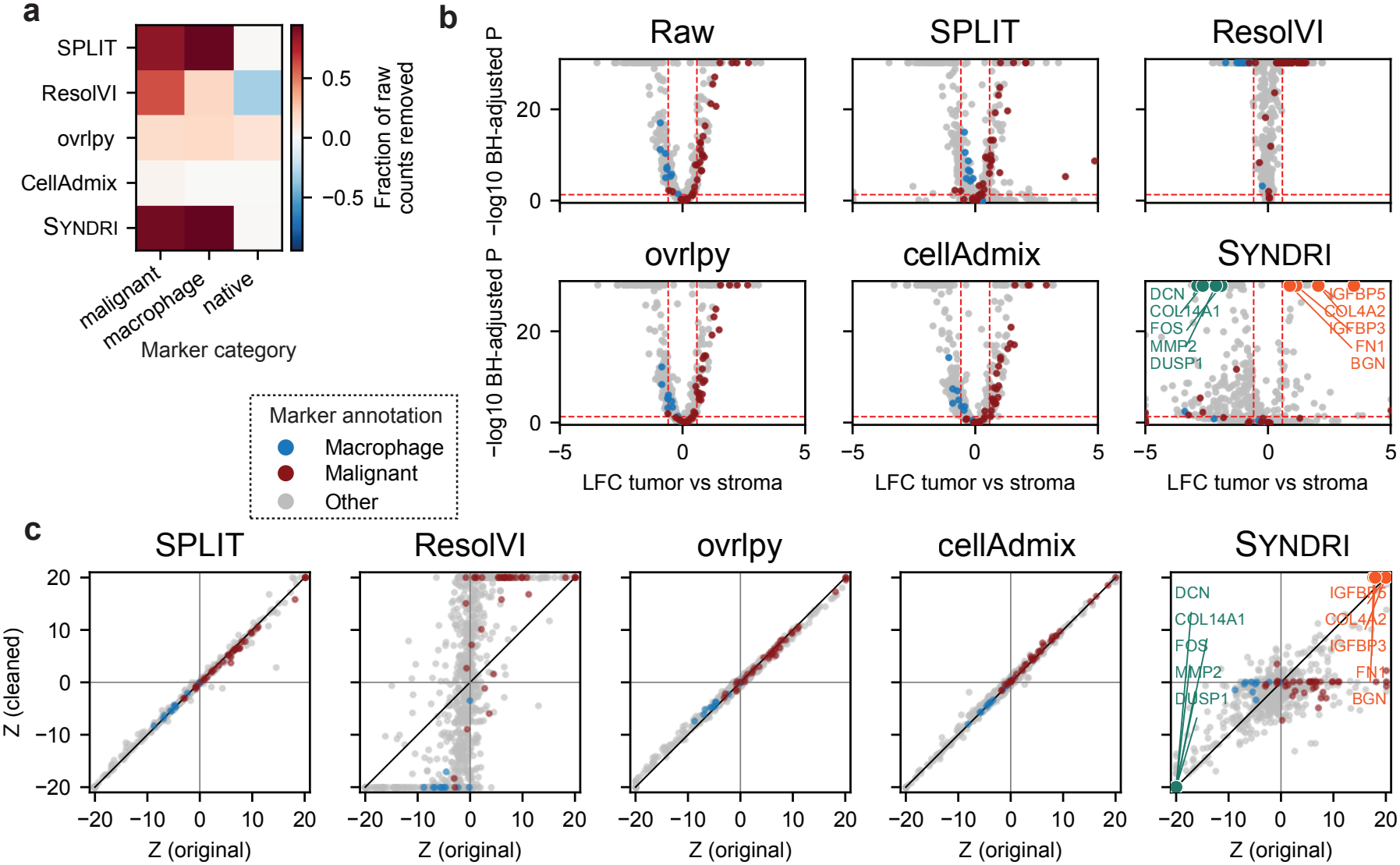
Syndri removes spillover-driven tumor and macrophage markers from differential expression in CosMx NSCLC fibroblasts. **a**. Fraction of raw transcript counts removed by each correction method for malignant, macrophage, and native fibroblast marker sets in annotated fibroblasts. **b**. Volcano plots of differential expression between fibroblasts in tumor and stromal regions, pooled across three replicates from one donor, before correction (Raw) and after correction by each method. Selected stromal- and tumor-associated fibroblast genes are highlighted in green and orange, respectively. **c**. Comparison of gene-wise Wilcoxon Z statistics for tumor-versus-stroma differential expression before and after correction. The diagonal denotes unchanged differential signal and movement toward Z = 0 indicates attenuation of the original differential expression. The same genes from **b** are highlighted for Syndri.

Following the fibroblast differential-expression analysis introduced in cellAdmix [12], we compared fibroblasts located in tumor-associated and stromal regions of the NSCLC dataset (**Fig**. 4). To assess the removal of potentially contaminating expression, we used malignant and macrophage marker sets derived from the cellAdmix analysis that are minimally expressed in fibroblasts in the matched scRNA-seq reference. A separate set of native fibroblast markers was used to assess preservation of endogenous expression. Differential expression between tumor-associated and stromal fibroblasts was then evaluated independently for each correction method using a standardized Wilcoxon rank-sum procedure with Benjamini–Hochberg adjustment (**Supplementary Methods** A.7).

We first quantified how strongly each correction method removed transcripts belonging to these three marker sets by calculating the fraction of transcripts present in the uncorrected data that were removed after correction (**Fig**. 4a). Syndri removed nearly all malignant-marker transcripts and a similarly large fraction of macrophage-marker transcripts, while removing very little native fibroblast-marker expression. SPLIT also removed substantial malignant and macrophage signal, but less completely than Syndri. In contrast, ovrlpy and cellAdmix removed little of the malignant-marker signal, and ResolVI produced reconstructed expression values instead of a directly corrected observed count matrix, complicating interpretation of transcript removal. Although cellAdmix was originally evaluated using this fibroblast differential-expression task, our rerun using the recommended settings (“cell-bridging” and factors = 1.5× number of cell types) showed substantially less removal of malignant- and macrophage-associated expression than reported in its original study. Differences in analysis procedures and the incomplete availability of one replicate may contribute to this discrepancy.

These differences changed the DE conclusions. In the uncorrected data, many significant DEGs were malignant or macrophage markers, reflecting neighboring tumor and immune cells rather than the fibroblast program (**Fig**. 4b). Syndri was the only method after which nearly all of these markers became non-significant; they remained among the DEGs after SPLIT, ovrlpy, and cellAdmix. ResolVI produced a substantially different DE landscape, likely because its posterior reconstruction assigns small fractional values to many genes, including genes not detected in the original data. Comparing gene-wise Wilcoxon *Z* statistics before and after correction showed the same pattern (**Fig**. 4c): for SPLIT, ovrlpy, and cellAdmix, most genes, including most malignant and macrophage markers, stayed near the diagonal, whereas Syndri moved nearly all malignant and macrophage markers toward *Z* = 0. Many non-marker genes retained substantial DE signal under Syndri.

The strongest DEGs remaining after Syndri correction were associated with fibroblast state and matrix organization. Tumor-region fibroblasts were enriched for *IGFBP5, COL4A2, IGFBP3, FN1*, and *BGN*, and stromal fibroblasts for *DCN, COL14A1, FOS, MMP2*, and *DUSP1*. These are consistent with the heterogeneous fibroblast states described across tissues and in cancer [24], including adventitial, *COL14A1*-associated matrix-fibroblast programs in human NSCLC [25] and murine lung [26]. Removing malignant and macrophage signal therefore did not collapse biological variation among fibroblasts.

Together, these results show that spillover distorts spatial DE analysis and that methods differ greatly in their ability to remove it. Syndri removed malignant and macrophage signal from fibroblasts while preserving native fibroblast markers and plausible fibroblast-state differences, improving the reliability of downstream analyses such as marker discovery and tumor–stroma comparisons.

### Syndri responds to cellular context

We tested whether Syndri uses a cell’s neighborhood, beyond the cell’s own expression, to decide which transcripts to remove. In the breast cancer Xenium dataset, we held each query cell’s expression profile fixed and replaced the cell types of its five neighbors with those from other observed neighborhoods, varying the number of malignant neighbors (**Fig**. 5a). Any change in the model’s output is therefore attributable to neighborhood context alone.

**Figure 5:**
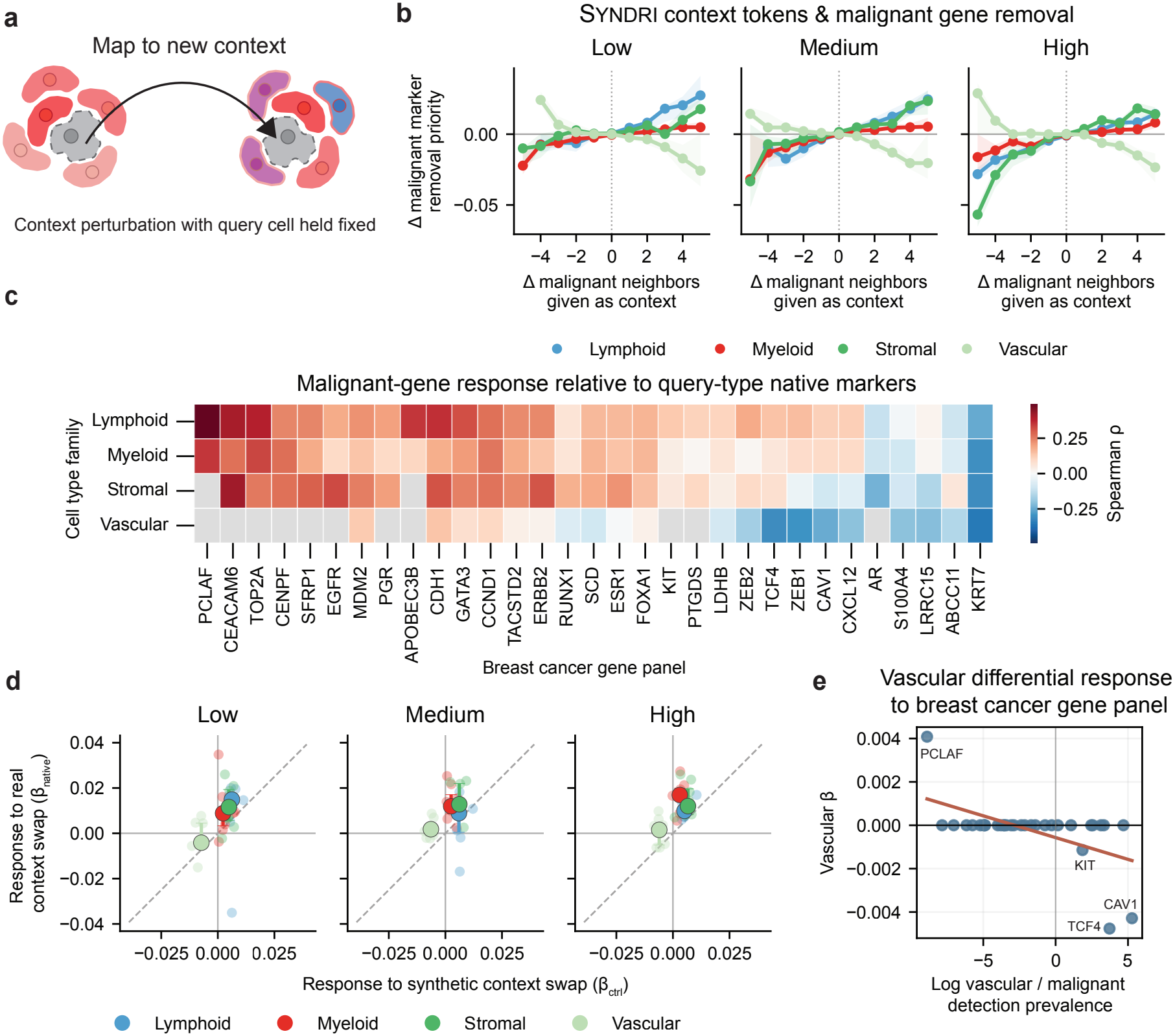
Neighborhood context changes Syndri’s spillover predictions in a cell-class- and gene-specific manner. **a**. Context-remapping experiment. Each query cell’s expression profile and cell type are held fixed while the cell types of its five neighbors are replaced with those from other observed neighborhoods. **b**. Change in malignant-marker removal priority (AUROC of malignant markers versus markers of the query cell’s own class) as the number of malignant neighbors increases, stratified by cell class and baseline malignant-marker expression. Lines show medians across tissue sections; shaded regions show interquartile ranges. **c**. Gene-specific context response for the 32-gene breast cancer marker panel, grouped by query-cell family. Colors indicate the median Spearman correlation, across supported tissue sections and query strata, between changes in malignant-neighbor count and changes in each gene’s removal priority relative to native markers. Gray entries indicate insufficient support or undefined correlations. **d**. Context-response slopes from the remapping experiment compared with the association between malignant-neighbor count and removal priority in unperturbed tissue, for each cell class and baseline expression level. **e**. In vascular cells, context-response slope of each malignant-panel gene versus its prevalence in capillary endothelial relative to malignant cells in the matched single-cell reference.

We summarized the output as the malignant-marker removal priority: the AUROC with which Syn-dri ranks a 32-gene breast cancer marker panel above markers of the query cell’s own cell class for removal. We grouped query cells by cell class (lymphoid, myeloid, stromal, or vascular) and by baseline malignant-marker expression (low, medium, or high). Adding malignant neighbors increased the removal priority of malignant markers in lymphoid and myeloid cells, and in stromal cells with medium or high baseline expression, but decreased it in vascular cells (**Fig**. 5b). These effects were reproducible across tissue sections. Neighborhood context therefore shifts Syndri’s predictions in a cell-class-dependent manner rather than uniformly raising the probability that malignant markers are removed.

The effect also varied by gene (**Fig**. 5c). *CDH1, GATA3, CCND1, TACSTD2*, and *ERBB2* showed positive responses across several cell classes, whereas other genes showed weak, negative, or class-specific responses. In vascular cells, genes that were more prevalent in capillary endothelial than in malignant cells of the matched single-cell reference showed more negative responses (Spearman *ρ* = *−*0.52, two-sided permutation *P* = 6 × 10^*−*4^, *n* = 32 genes; **Fig**. 5e), for example *CAV1*. Syndri thus appears to avoid removing transcripts that are plausibly endogenous to the query cell, so context adjusts removal priority gene by gene rather than acting as a global malignant-context multiplier.

Finally, these perturbation effects agreed with unperturbed tissue (**Fig**. 5d). In most combinations of cell class and baseline expression, the response to perturbation matched in direction the association between malignant-neighbor count and removal priority in the original tissue, most strongly at high baseline expression; vascular cells showed weaker agreement in some strata. Together, these results show that Syndri uses neighborhood context to adjust which transcripts are removed in a manner specific to cell class and gene, and consistent with patterns in native tissue.

## Discussion

We developed Syndri, a framework that corrects transcript spillover in imaging-based spatial transcriptomics by casting it as a supervised learning problem. Syndri generates labeled training data by assigning spillover-free scRNA-seq profiles to the cell positions of an ST sample and simulating contamination between neighboring cells, so that the true expression of every cell is known. A transformer trained on these data uses each cell’s expression, cell type, and neighborhood to predict an admixture probability for every observed gene, and Syndri removes genes predicted to be contaminated. This design lets Syndri operate directly on cell-by-gene count matrices, without transcript coordinates or deconvolution.

Across MERFISH, CosMx, and Xenium datasets, Syndri outperformed existing methods in cell-type separation, reference similarity, and heterogeneity preservation, improving cell-type separation without sacrificing within-type variation. In NSCLC, it removed spillover-driven tumor and macrophage markers from the DEGs between fibroblasts in tumor and stromal regions while retaining DEGs consistent with known fibroblast states. Controlled context-remapping experiments further showed that Syndri’s predictions change with neighborhood composition in a cell-class- and gene-specific manner.

Several limitations remain. First, Syndri relies on synthetic supervision, and its performance therefore depends on how well simulated admixture approximates the mechanisms underlying transcript spillover in experimental data. Although Syndri-Sim reproduces several features of real spatial expression and provides known contamination labels for benchmarking, it does not fully recapitulate the observed patterns of cross-cell-type marker co-detection. Moreover, because the simulated benchmark uses the same general contamination mechanism as training, performance on synthetic data may not fully reflect generalization to experimental spillover. Second, our training procedure introduces contamination only at genes absent from the native query cell, whereas real spillover can also increase counts of endogenously expressed genes. Consequently, Syndri’s whole-gene correction strategy cannot distinguish endogenous and contaminating transcripts of the same gene and may remove genuine expression when both are present. Finally, Syndri requires cell-type annotations for query cells and their neighbors, making its predictions potentially sensitive to annotation errors, particularly in heterogeneous tissues or regions containing transcriptionally similar cell populations.

These limitations suggest several directions for future development. More realistic synthetic training data could incorporate gene-specific contamination rates, segmentation geometry, transcript density, and three-dimensional tissue organization to better approximate experimental spillover. Extending Syndri from binary gene-level detection to estimation of contaminating transcript counts could enable partial correction of genes containing both endogenous and misassigned expression. Neighborhood representations could also incorporate continuous spatial distances, cell morphology, or graph-based tissue structure rather than relying exclusively on categorical cell-type tokens, potentially reducing dependence on accurate cell-type annotations. Finally, adaptive correction strategies that account for uncertainty in predicted contamination could replace the current fixed removal thresholds, allowing correction strength to vary across cells and tissue environments. Together, these extensions could improve the robustness and applicability of supervised spillover correction across diverse spatial transcriptomics datasets.

In general, Syndri provides a flexible tool for removing spillover while preserving biological variation, supporting more reliable analysis of cellular organization and gene-expression programs across spatial technologies and tissues.

## Methods

### Synthetic admixture generation

The absence of experimentally measured ground-truth transcript spillover impedes direct supervised training on spatial transcriptomics data. We therefore construct synthetic training examples from scRNA-seq profiles, for which the origin of artificially introduced transcripts is known by construction. Each training example consists of a query cell, a set of neighboring source cells, a synthetically admixed expression profile, and binary gene-level labels indicating which genes received expression from the source cells.

#### Construction of pseudo-spatial single-cell data

Because scRNA-seq measurements lack spatial coordinates, we construct a pseudo-spatial version of each single-cell sample before generating synthetic admixture, where we define admixture as the result of adding spillover to a cell’s endogenous expression. For each dataset, we first select a representative ST slice whose cell-type adjacency distribution is most similar to the other slices in the dataset. To find this slice, we build a spatial KNN graph over the cells for each ST slice, and calculate the adjacency-enrichment profiles based on observed vs expected neighbor fractions computed from the cell type abundance for each directed cell-type pair. We use the log of the observed vs expected ratio for each pair as an adjacency-enrichment profile and select the slice with maximum mean Spearman correlation of its profile with every other slice.

The selected spatial slice and corresponding scRNA-seq reference are jointly embedded using Harmony [27]. Briefly, the two datasets are concatenated, library-size normalized, log-transformed, scaled, and projected into a 30-dimensional PCA space. Harmony is then applied using data source (spatial or single-cell) as the batch covariate, yielding a shared representation for the spatial and single-cell profiles.

To transfer the observed tissue organization to the single-cell reference, we assign an scRNA-seq profile to each unique spatial-cell coordinate in the representative slice. For spatial cell *r*, let 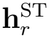 denote its Harmony embedding and let 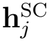 denote the embedding of single-cell reference cell *j*. We first identify the *M* = 50 nearest single-cell candidates to each spatial cell using Euclidean distance in the Harmony embedding,

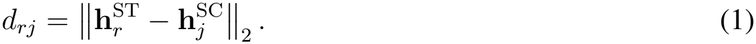

Candidate matches are restricted to those with the same cell-type annotation as the spatial cell and are converted to distance-based sampling weights,

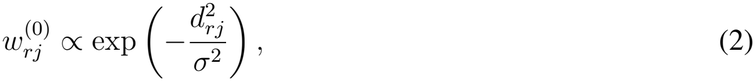

where *σ* is the median candidate-match distance across the mapping. To discourage a small number of reference cells from being repeatedly assigned throughout the pseudo-spatial tissue, these weights are further adjusted by a reuse penalty. If *c*_*j*_ denotes the frequency with which reference cell *j* occurs as the nearest candidate across spatial positions, its sampling weight is proportional to

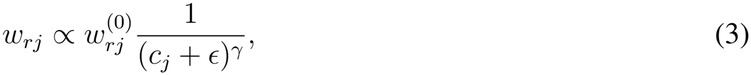

where *γ* controls the strength of the reuse penalty. The adjusted weights are normalized across the candidate cells for each spatial position, and one scRNA-seq cell is sampled from the resulting distribution. *γ* = 9 and a hard max reuse cap of 5000 per sample were used for all datasets.

The resulting pseudo-spatial dataset contains one sampled scRNA-seq expression profile for each unique spatial coordinate of the representative slice, and individual reference profiles may therefore be reused at multiple spatial positions. This procedure preserves the experimentally observed arrangement of cell types and local tissue neighborhoods while retaining scRNA-seq expression profiles as the substrate for synthetic admixture generation. Synthetic source cells – those from which synthetic spillover originates – are subsequently defined using physical proximity in this transferred coordinate system.

#### Sampling of Admixed Transcripts

For each query cell, we construct a candidate contaminating expression profile by aggregating the expression counts of its five nearest eligible donor cells with different annotated cell types. Let

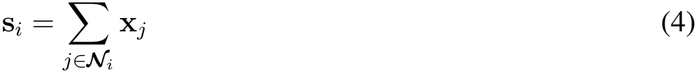

denote the aggregate expression of the source-cell neighborhood.

To make synthetic contamination identifiable at the gene level, candidate admixed genes are restricted to genes not already detected in the query cell 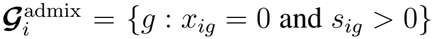 . Thus, any nonzero expression introduced at an eligible gene can be attributed unambiguously to synthetic admixture.

For each training example, a target final admixture fraction *τ* was sampled uniformly from [0.7, 0.85]. This specifies the desired fraction of the final expression profile attributable to introduced transcripts, although the realized fraction may differ because donor expression is finite and whole-gene counts are added without truncation.

The target number of contaminating transcripts is therefore

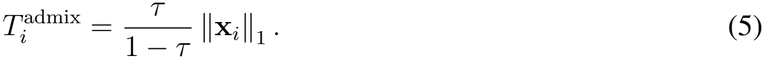

Sampling was skipped when this target was below the configured minimum of 10 transcripts. Otherwise, if the available eligible donor counts were insufficient to reach the target, all eligible pooled counts were added. When sufficient counts were available, eligible genes were randomly permuted and their entire pooled counts were added until the target was reached or exceeded. Consequently, the realized admixture fraction could fall below the sampled target because of limited donor availability, or exceed it because genes were transferred as whole count blocks.

#### Gene-Level Admixture Labels

Because the synthetic admixture procedure explicitly records which genes receive contaminating transcripts, each training example is paired with a binary gene-level target. For query cell *i* and gene *g, y*_*ig*_ = I[*a*_*ig*_ > 0], where *a*_*ig*_ is the total number of synthetic contaminating transcripts added for gene *g*. Thus, *y*_*ig*_ = 1 indicates that the observed expression of gene *g* contains synthetically introduced signal, while *y*_*ig*_ = 0 indicates that no synthetic admixture was added for that gene. These labels provide direct supervision for the gene-level admixture prediction task used to train Syndri.

### Syndri Architecture

Admixture cannot always be identified from the expression of an individual gene in isolation. Whether expression is plausible depends on the identity of the query cell, its complete expression profile, and the types of cells present in its local spatial environment. We use a transformer architecture previously developed by our group that jointly represents gene expression and cellular context [28].

#### Gene Identity and Expression Tokenization

For each query cell, the input contains one token for each gene with nonzero observed expression. Gene tokens are ordered by decreasing expression, as introduced in Geneformer [29]. Each token combines a learned embedding of the gene identity with a learned linear projection of its expression count, where the counts are neither normalized nor log-transformed.

#### Cell-Type and Neighborhood Tokenization

The gene tokens are preceded by metadata tokens describing cellular context. For each query cell, we include one learned cell-type token for each of its five nearest spatial neighbors with a different annotated cell type, followed by a learned token representing the query cell’s own annotated type. Thus, the input represents the query expression profile together with the types of the cells in its local neighborhood and the query cell’s own type. Finally, we prepend a `[CLS]` token to the complete sequence. For the separate Syndri-Sim benchmark model, the neighborhood representation was expanded to nine neighbor cell-type tokens to match the nine-donor simulation configuration.

#### Transformer Encoder and Prediction Head

The input embeddings are augmented with learned, BERT-style positional embeddings and processed by a four-layer transformer encoder with eight self-attention heads and a hidden dimension of 128 [30]. Each layer uses a 512-dimensional feed-forward sublayer with GELU activation; dropout is 0.1.

For admixture prediction, a shared linear head is applied to each contextualized gene-token embedding. The `[CLS]` token and cell-type prefix-token embeddings (six in the primary models and ten in the Syndri-Sim benchmark model) are excluded from gene-level prediction. For query cell *i* and gene *g*, let **h**_*ig*_ ∈ ℝ ^128^ denote the contextualized gene-token representation. The head maps each gene embedding to a scalar admixture logit, *z*_*ig*_ = **w**^⊤^**h**_*ig*_ + *b*, from which the gene-level admixture probability is obtained as *p*_*ig*_ = *σ* (*z*_*ig*_).

Although predictions are produced separately for each gene token, each **h**_*ig*_ incorporates information from the complete input sequence through transformer self-attention [31]. Predictions can therefore depend on the query expression profile, its cell-type annotation, and the cell types in its spatial neighborhood.

### Admixture Correction

After inferring gene-level admixture probabilities with a trained Syndri model, we construct corrected expression matrices by zeroing observed cell–gene counts predicted to contain contamination. Correction thresholds are determined separately for each annotated cell type within each spatial slice. Within each cell-type group, expressed cell–gene entries with predicted admixture probability at least 0.5 are ranked in descending probability order. Entries are selected for removal until the cumulative number of removed transcripts reaches a target fraction (*ρ* = 0.5) of the group’s total observed transcript counts, or until no eligible entries remain. All entries at or above the resulting probability threshold are set to zero.

Because removal operates on complete cell–gene counts, and multiple entries may share the threshold probability, the achieved removal fraction may differ from the target. Counts associated with probabilities below 0.5 are always retained. The target removal fraction is a target rather than a fixed amount of expression removed; when the cumulative counts associated with predictions above the minimum probability threshold are insufficient to reach the target, fewer transcripts are removed.

For evaluation on simulated ST data with known contamination labels (**Fig**. 3), we separately assessed continuous admixture scores using AUPRC and binarized predictions using F1-score. Binary operating points were selected using thresholds that maximized F1-score against the known contamination labels. These oracle thresholds were used solely for retrospective benchmarking and not for correction of experimental ST data.

### Spillover correction metrics

We evaluated spillover-correction methods along three complementary dimensions: cell-type separation, similarity to the matched scRNA-seq reference, and preservation of within-cell-type heterogeneity. Cell-type separation was quantified using adjusted Rand index (ARI), normalized mutual information (NMI), and silhouette score; ARI and NMI were computed using `scib-metrics` [21, 32] and silhouette score was computed using the `scikit-learn` implementation [33]. Reference similarity was measured using cosine similarity between cell-type pseudobulk profiles, and heterogeneity preservation was assessed by comparing gene-wise expression variance between the spatial data and the matched scRNA-seq reference.

#### Reference similarity

To quantify agreement between corrected spatial expression profiles and the matched scRNA-seq reference, we constructed cell-type pseudobulk profiles by summing counts across all cells of each annotated type. Spatial and reference pseudobulks were independently normalized to 10,000 counts and transformed using log(1 + *x*). For method *m*, slice *s*, and cell type *c*, cosine similarity across shared genes was calculated as

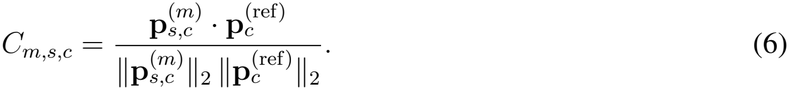

The slice-level reference-similarity score was the arithmetic mean across matched cell types,

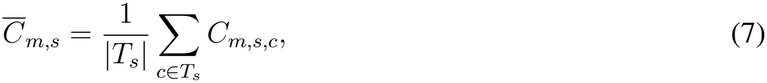

where *T*_*s*_ denotes the set of matched cell types used for slice *s*. The same set of matched cell types was used across methods for a given slice, so each cell type contributed equally to the slice-level score.

#### Preservation of within-cell-type heterogeneity

To assess whether correction preserved expression variation within annotated cell types, we compared gene-wise variance in the corrected spatial data with that of the matched scRNA-seq reference. Within each slice and cell type, counts were normalized to 10,000 per cell and transformed using log(1 + *x*). We retained shared genes with nonzero variance in the reference that were detected in at least 5% of reference cells. Comparisons were restricted to cell types represented by at least 50 spatial cells and 50 reference cells and containing at least two eligible genes.

For method *m*, slice *s*, and cell type *c*, we calculated

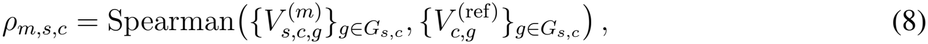

Where 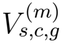 and 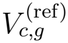 denote the across-cell variance of normalized log expression for gene *g* in the spatial and reference data, respectively, and *G*_*s,c*_ is the set of eligible shared genes. The slice-level heterogeneity-preservation score was the median *ρ*_*m,s,c*_ across eligible cell types. Higher values indicate better preservation of the relative pattern of gene-wise variation observed in the reference.

#### Cross-metric relative-rank summary

To summarize performance across metrics with different numerical scales, methods were ranked separately within each slice for each of the five metrics above, treating higher values as better. If method *m* had rank *r*_*m,s,k*_ among *M*_*s*_ evaluated methods for metric *k*, the rank was normalized as

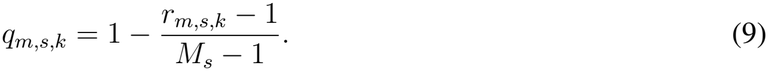

This transformation assigns a score of 1 to the best-performing method and 0 to the worst-performing method within each slice and metric.

Normalized metric scores were first averaged across slices within each dataset. Cell-type separation was defined as the equal-weight mean of ARI, NMI, and silhouette score. The overall score was then calculated as the equal-weight mean of cell-type separation, reference similarity, and heterogeneity preservation. Datasets were weighted equally in across-dataset summaries.

### Benchmarking with Syndri-Sim

We constructed Syndri-Sim to evaluate spillover correction against known native and added expression counts. Single-cell RNA-seq expression profiles were assigned to observed CosMx cell positions, after which synthetic spillover was introduced from neighboring cells. This construction separates the expression used to generate the benchmark from the real spatial measurements used to assess distributional similarity.

#### Construction of simulated spatial data

The NSCLC scRNA-seq reference was partitioned by patient into training, validation, and test sets comprising five, one, and one donors, respectively. Reference profiles from each partition were independently mapped to observed CosMx cell positions using Harmony integration and cell-type-matched nearest-neighbor sampling. Training, validation, and test sets were constructed using three, one, and four spatial slices, respectively. No reference cells were shared across partitions.

For each pseudo-spatial query cell *i*, we selected up to nine nearest neighboring donor cells of different annotated cell types. Let *X*_*ig*_ denote the native expression count of gene *g* in cell *i, t*_*j*_ the cell type of donor *j*, and ***N*** (*i*) the selected donor neighborhood.

To account for differences in the contribution of different donor cell types to observed contamination, donor expression was scaled by cell-type-specific weights 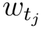, estimated as described below. Weighted donor counts were converted to integers by stochastic rounding:

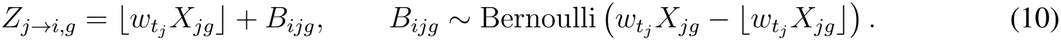

As in the original training procedure, synthetic contamination was restricted to genes absent from the native query. The eligible donor pool was therefore

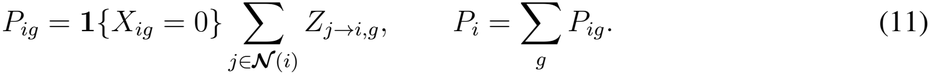

For each query, we sampled a transfer fraction and corresponding target number of added transcripts:

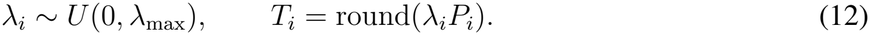

Genes with positive eligible donor counts were randomly permuted, and their entire pooled counts were added sequentially until the cumulative addition reached or exceeded *T*_*i*_. The final gene was not truncated. The resulting simulated expression was

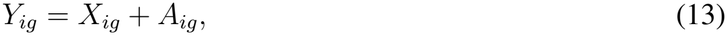

where *A*_*ig*_ denotes the added contaminating counts. Donor expression was always taken from the original native profiles, so contamination was not propagated between cells. Unlike the original training sampler, which specifies a target final contamination fraction *τ*, the parameter *λ*_*i*_ specifies a fraction of the eligible weighted donor pool.

#### Calibration of synthetic spillover

We calibrated donor-type contributions and the maximum transfer fraction using a held-out set of 3 spatial slices. Donor-type weights were estimated from the excess detection of cell-type-specific marker genes in real CosMx recipient cells relative to the matched scRNA-seq reference.

For donor type *d*, let *M*_*d*_ denote its marker set, 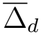 the mean excess marker-detection frequency across supported recipient types and calibration slices, and 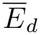 the corresponding mean availability of eligible native marker counts in neighboring donor cells. We defined the unnormalized donor contribution as

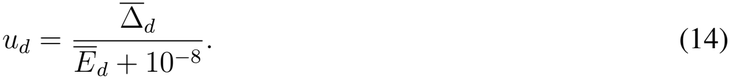

Weights were normalized to mean one across supported donor types and shrunk toward one:

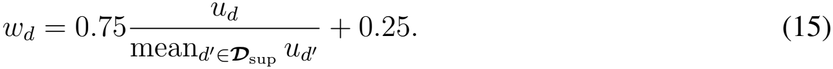

Unsupported donor types received *w*_*d*_ = 1. The resulting weights were fixed before test-set simulation and were not renormalized within individual neighborhoods. The marker-selection and support criteria are described in Supplementary Methods A.9.

We selected *λ*_max_ by comparing simulated and real cell-level expression distributions using squared maximum mean discrepancy (MMD^2^). We evaluated

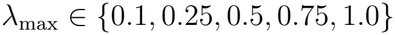

using three simulation seeds and up to 500 fixed query cells per supported cell type and calibration slice.

Expression vectors were normalized to 10,000 counts per cell and log-transformed. For simulated and real expression samples 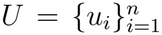 and 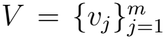, respectively, we used the biased MMD^2^ estimator:

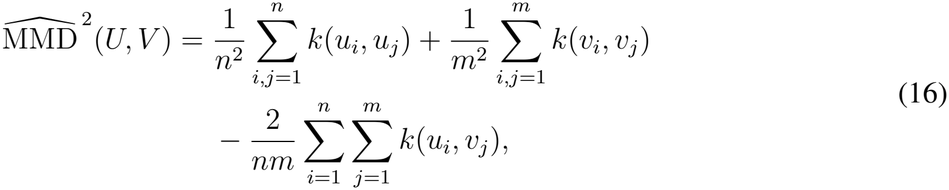

where the radial basis function kernel was

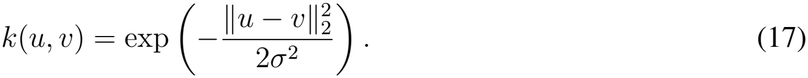

The bandwidth *σ*^2^ was determined from the median positive squared pairwise distance among pooled native pseudo-spatial and real expression profiles within each slice–cell-type stratum.

MMD^2^ values were averaged equally across supported cell types, calibration slices, and simulation seeds. This procedure selected *λ*_max_ = 0.25. Neither donor-type weights nor the transfer parameter were calibrated using held-out test slices.

#### Evaluation against known spillover

The calibrated simulator was applied to four held-out CosMx test slices. Syndri was retrained using the calibrated simulation procedure and the original training and validation partitions, with no reference cells shared with the test partition. SPLIT and ResolVI were evaluated using the same saved simulated expression matrices.

To evaluate methods requiring transcript coordinates, we additionally generated transcript-level positions within the observed cell segmentation masks. Native transcripts were sampled uniformly within the polygon of their corresponding recipient cell. Added transcripts were sampled within the recipient mask with a spatial bias toward the known donor cell. For an added transcript originating from donor j, the relative sampling weight at position x was

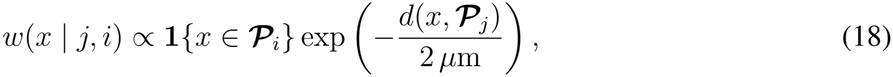

where ***P***_*i*_ and ***P***_*j*_ denote the recipient and donor segmentation regions, respectively, and *d*(*x*, ***P***_*j*_) is the distance from position *x* to the donor region. Discrete *z*-coordinates were sampled from the empirical slice-wide distribution. This procedure preserved the simulated transcript counts and known donor– recipient assignments.

The generated transcript coordinates enabled evaluation of ovrlpy and cellAdmix alongside Syn-dri, SPLIT, and ResolVI. All methods were evaluated using the same simulated expression counts and known contamination labels. Transcript-level methods additionally received the simulated transcript coordinates, with method-specific settings described in Supplementary Methods A.8. Because native and added counts were retained separately, the ground-truth contamination label for each expressed cell–gene entry was known:

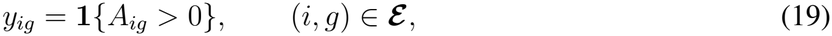

where

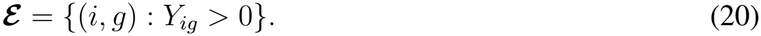

For Syndri, genes were ranked using their predicted admixture probabilities *p*_*ig*_. For comparison methods returning corrected expression counts *C*_*ig*_, we quantified the degree of removal as

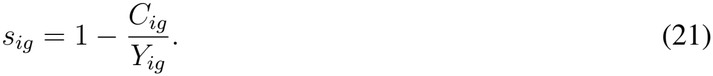

Larger scores indicate greater removal of the observed expression.

The binarized version of Syndri reported in **Fig**. 3d was constructed by first finding the oracle probability threshold that maximized the F1. Binarized Syndri simply treats expression removal as all or nothing depending on whether each gene is above the oracle threshold. For methods returning corrected counts, continuous removal scores were calculated as one minus the ratio of corrected counts to the synthetically admixed observed counts for each expressed cell–gene entry.

Predictions and labels were pooled across all four test slices, with each expressed cell–gene entry contributing equally. We computed precision–recall curves and their area (AUPRC) from the continuous removal scores. We also evaluated binarized predictions using F1-score,

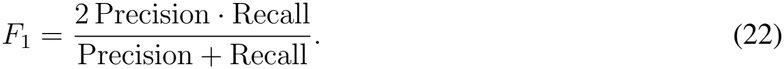

For comparison methods, thresholds maximizing F1-score were determined using the simulated ground-truth labels. These oracle thresholds were used only for retrospective benchmarking and do not represent independently validated deployment settings. The full evaluation procedure is described in Supplementary Methods A.8.

### Neighborhood context-response analysis

#### Context remapping

To assess whether neighborhood composition influences Syndri’s predictions independently of query-cell expression, we performed a context-remapping experiment on eight breast cancer Xenium sections. For each exact cell type and section, we sampled up to 300 query cells and evaluated their expression profiles under alternative neighborhood contexts drawn from other cells of the same type and section. Each query’s expression and cell-type token were held fixed while the five neighbor cell-type tokens were replaced, yielding up to 30 alternative contexts spanning different numbers of malignant neighbors.

Let 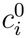 denote the original neighborhood context of query cell *i*, and *c* an alternative context. We define *m*(*c*) as the number of malignant-cell tokens among the five neighbors. The change in malignant-neighbor count under context substitution is

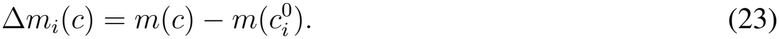

Because query expression and identity were unchanged, differences in predicted admixture probabilities between 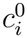 and *c* reflect the model’s response to the supplied neighborhood context.

#### Marker removal priority

We quantified the relative removal priority of malignant markers using the predicted admixture probabilities of a breast cancer panel compared with native markers of the query cell type. For query *i*, let *M*_*i*_ and *N*_*i*_ denote the sets of expressed malignant-panel and native-marker genes, respectively, and let *p*_*ig*_(*c*) be the predicted admixture probability for gene *g* under context *c*.

We defined malignant-marker removal priority as the AUROC comparing these two gene sets:

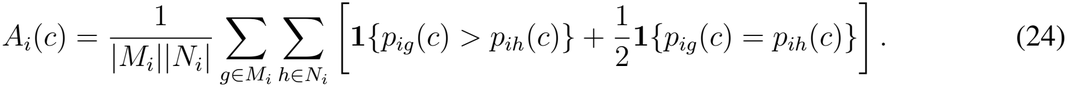

Larger values indicate that malignant-panel genes are ranked more highly for removal than the query cell’s native markers. This statistic measures relative removal priority rather than the fraction of transcripts removed.

For each context substitution, we calculated

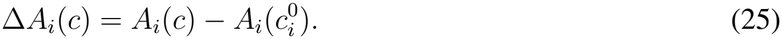

Positive values indicate increased malignant-marker removal priority relative to the original context.

Responses were grouped by cell-type family (lymphoid, myeloid, stromal, or vascular) and baseline malignant-marker burden. The latter was calculated from the original expression profile as

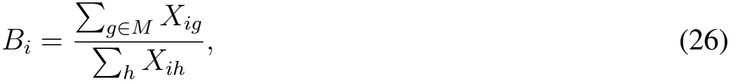

where *M* is the complete 32-gene panel and *X*_*ig*_ is the observed expression count. Queries were divided into low, medium, and high burden groups using tertiles calculated within each section and exact cell type. Sampling and aggregation additionally accounted for the original number of malignant neighbors, as described in Supplementary Methods A.9.

### Quantification of context response

We first examined changes in malignant-marker removal priority as a function of Δ*m*_*i*_(*c*), aggregating responses across query cells, cell types, and tissue sections.

To assess gene-specific responses, we defined the removal priority of an individual malignant-panel gene *g* relative to the query’s expressed native markers:

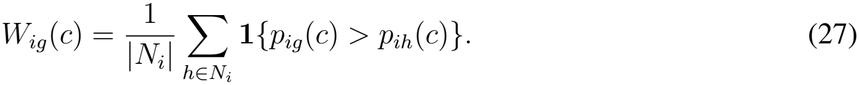

We then calculated

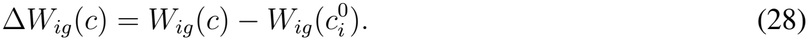

Gene-specific context response was measured using the Spearman correlation between Δ*m*_*i*_(*c*) and Δ*W*_*ig*_(*c*), calculated within section, cell-type family, and query stratum. Positive correlations indicate that increasing the number of malignant neighbors raises a gene’s removal priority relative to native markers.

We also compared the response to controlled context substitution with the association observed in unperturbed tissue. For each fixed query, we fitted

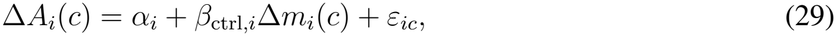

where *β*_ctrl,*i*_ measures the change in removal priority per additional malignant-neighbor token while query expression and identity remain fixed.

For the corresponding unperturbed queries, we fitted a weighted regression within each section *s*, exact cell type *t*, and baseline burden group *b*:

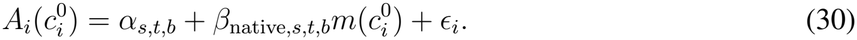

Here, *β*_native_ describes the association between malignant-neighbor count and removal priority across different cells in their original spatial contexts. Comparing the signs and magnitudes of *β*_ctrl_ and *β*_native_ allowed us to assess whether controlled changes in model input were consistent with associations observed in the original tissue. Unlike *β*_ctrl_, *β*_native_ is observational and may reflect other differences between cells.

Finally, we examined whether gene-specific responses in vascular cells were related to the plausibility of endogenous expression. For each malignant-panel gene, we estimated a context-response slope from *W*_*ig*_(*c*) and Δ*m*_*i*_(*c*) across eligible capillary endothelial queries. We compared the median gene-specific slope with its relative detection prevalence in capillary endothelial and malignant cells of the matched scRNA-seq reference:

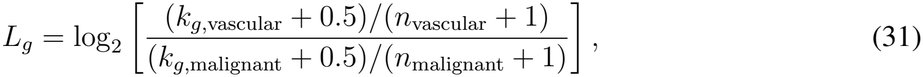

where *k*_*g,t*_ is the number of reference cells of type *t* expressing gene *g*, and *n*_*t*_ is the total number of reference cells of that type. Positive values indicate higher detection prevalence in vascular cells, whereas negative values indicate higher prevalence in malignant cells.

Detailed query selection, context sampling, statistical aggregation, and minimum-support criteria are provided in Supplementary Methods A.9.

## Acknowledgements

The authors thank Nicholas Ho, Shike Wang, Evan Trop, Fred Kolling, Aidan Zhang, Ana Prieto, and the rest of the Ma lab for their helpful discussions. This work was supported, in part, by the National Institutes of Health grants UM1HG011593 (J.M.), R03OD039980 (J.M.), UH3CA268202 (J.M.), R01HG012303 (J.M.), R01HG014954 (J.M.), R21DA061481 (J.M.), and U24HG012070 (J.M.). J.M. was additionally supported by the Ray and Stephanie Lane Professorship, a Guggenheim Fellowship from the John Simon Guggenheim Memorial Foundation, and a Google Research Award. S.K. was a Lane Fellow for part of the duration of the project. The funders had no role in study design, data collection and analysis, decision to publish or preparation of the manuscript.

## Code Availability

The source code for Syndri can be accessed at: https://github.com/gkrieg/Syndri

## Author Contributions

Conceptualization J.W., S.K.; Methodology J.W., S.K.; Software J.W., S.K.; Investigation J.W., J.M., S.K.; Writing J.W., J.M., S.K.; Funding Acquisition, J.M.

