## Supplementary Methods for "Syndri: Context-aware removal of transcript spillover in imaging-based spatial transcriptomics"

### A Supplementary Methods

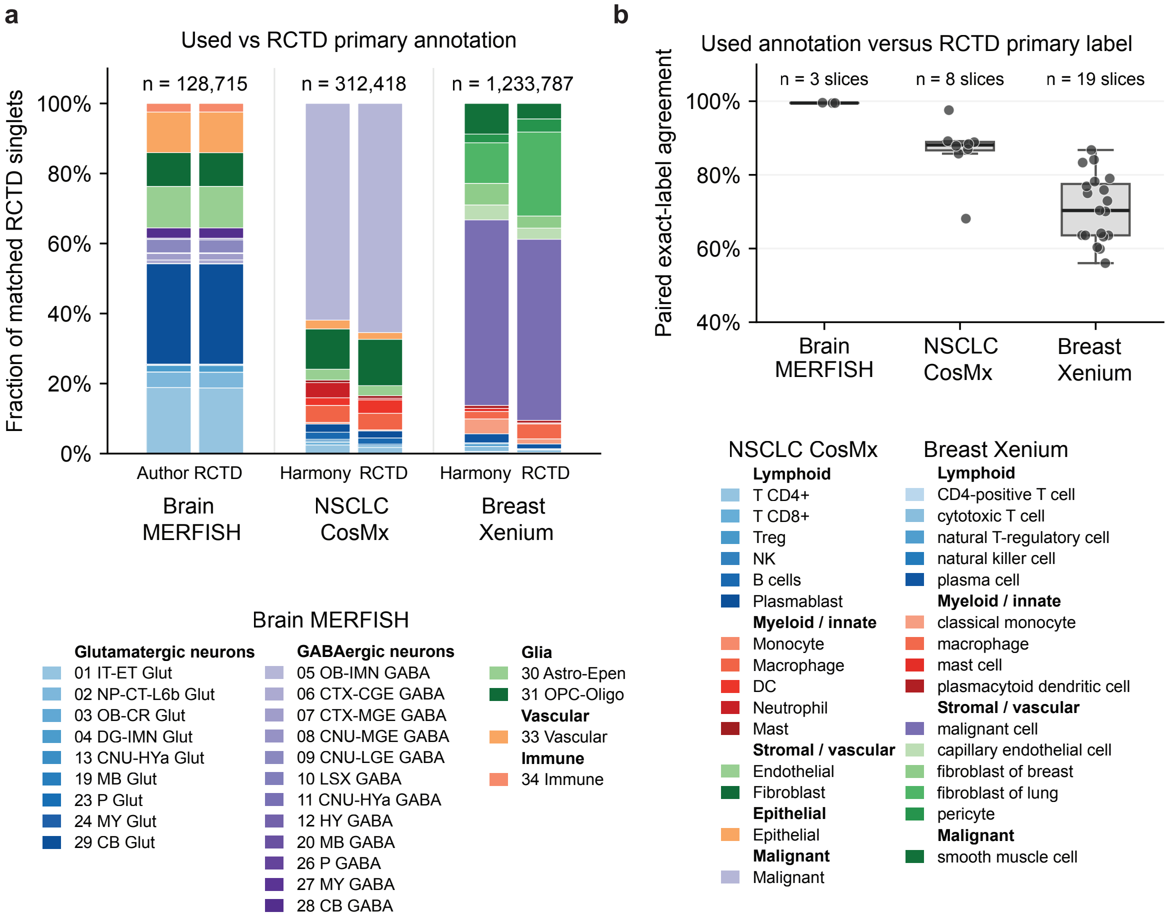


**Figure S1: Comparison of RCTD-derived and original cell-type annotations. a**. Cell-type composition of the annotations used in this study (Used) and the primary labels assigned by RCTD (RCTD) for the whole-mouse-brain MERFISH, NSCLC CosMx, and breast-cancer Xenium datasets. Bars show the proportion of each population annotated as each cell type, sorted by cell type family. Colors denote dataset-specific cell-type categories. **b**. Per-slice exact-label agreement between the annotations used in this study and the corresponding RCTD primary labels. Points represent individual spatial slices; boxplots summarize the distribution across slices.

#### A.1 RCTD annotations and SPLIT

SPLIT [1] uses RCTD [2] to estimate cell-type mixture proportions from a matched scRNA-seq reference, which are subsequently used to correct observed transcript counts. However, reference-based cell-type assignments can differ substantially across annotation methods. In a benchmark of reference-based annotation methods on breast cancer Xenium data, Cheng et al. [3] reported only 69.9% and approximately 55% agreement between RCTD-derived labels and manually curated annotations across two samples, respectively, demonstrating that substantial discrepancies can arise even when using a matched single-cell reference.

In contrast to SPLIT, SYNDRI uses existing cell-type annotations without requiring deconvolution. To characterize differences between the annotations used by these approaches, we compared the cell-type labels used in our analyses with the primary cell-type assignments obtained from RCTD (**Fig.** S1). The NSCLC CosMx and breast cancer Xenium datasets were annotated using the Harmony-based nearest-neighbor label-transfer procedure described in Section A.4, whereas the whole-mouse-brain MERFISH dataset retained the annotations provided by the original authors [4]. Comparisons were restricted to cells classified as singlets by RCTD. Agreement was nearly complete in the MERFISH dataset but lower and more variable in the CosMx and Xenium datasets, particularly the latter. These results demonstrate that SPLIT's deconvolution-based correction can depend on cell-type assignments that differ substantially from those used by SYNDRI, although this comparison does not independently establish which annotations are more accurate.

### A.2 Dataset details

All datasets used in this study are publicly available. The non-small cell lung cancer (NSCLC) CosMx Spatial Molecular Imager dataset was obtained from the NanoString public data portal (https://nanostring.com/products/cosmx-spatial-molecular-imager/ffpe-dataset/nsclc-ffpe-dataset/) [5]. The corresponding single-cell RNA-seq reference dataset is available from the Gene Expression Omnibus (GEO) under accession GSE127465 (https://www.ncbi.nlm.nih.gov/geo/query/acc.cgi?acc=GSE127465) [6].

The breast cancer Xenium dataset is available from GEO under accession GSE311609 (https://www.ncbi.nlm.nih.gov/geo/query/acc.cgi?acc=GSE311609) [1]. The corresponding single-cell RNA-seq reference dataset was obtained from the CELLxGENE Census collection (https://cellxgene.cziscience.com/collections/bd552f76-1f1b-43a3-b9ee-0aace57e90d6) [1].

The whole-mouse-brain MERFISH dataset was obtained from the CELLxGENE Census collection (https://cellxgene.cziscience.com/collections/0cca8620-8dee-45d0-aef5-23f032a5cf09) [4]; the non-imputed MERFISH measurements were used in this study. The corresponding single-cell RNA-seq reference data were obtained from the Allen Brain Cell Atlas whole-mouse-brain 10x Genomics dataset (https://github.com/AllenInstitute/abc_atlas_access/blob/main/descriptions/WMB-10X.md) [7].

### A.3 Implementation Details of Comparison Methods

#### *A.3.1 SPLIT*

Spatial transcriptomics data was re-annotated for cell-types using RCTD and the corresponding scRNA-seq reference. SPLIT was run in "doublet-mode".

#### *A.3.2 ResolVI*

ResolVI from scvi-tools [8] version 1.3.3 was run in semi-supervised mode, trained per slice, and then corrected counts were sampled from the updated model's posterior [9]. A maximum of 100 epochs were used for each training run. The posterior was sampled 3 times per cell, then the median of the samples was taken to obtain the corrected expression profile.

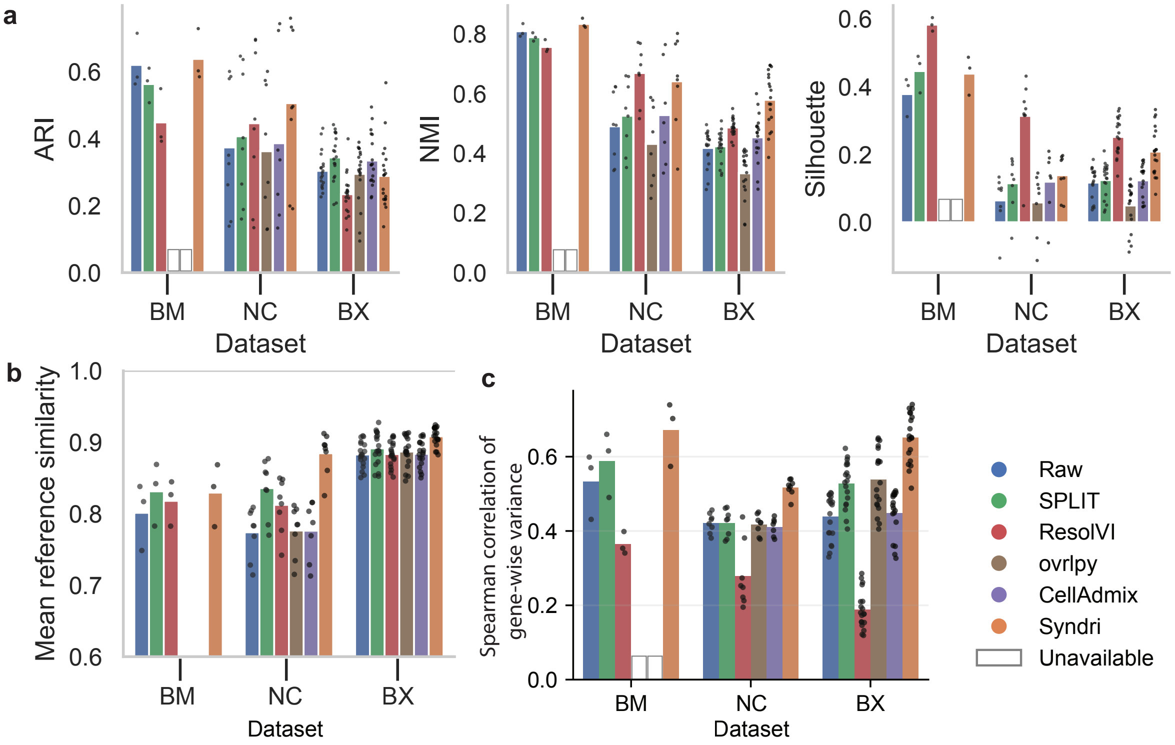


**Figure S2: Evaluation metrics for Syndri and baseline spillover-correction methods. a**. Cell-type separation measured by adjusted Rand index (ARI), normalized mutual information (NMI), and silhouette score for whole-mouse-brain MERFISH (BM), NSCLC CosMx (NC), and breast-cancer Xenium (BX). **b**. Mean cosine similarity between cell-type pseudobulk expression profiles in each spatial slice and the matched scRNA-seq reference. **c**. Preservation of within-cell-type heterogeneity, measured as the Spearman correlation between gene-wise expression variance in each corrected spatial dataset and the matched scRNA-seq reference. Bars show the mean across slices within each dataset, and points indicate individual slices. White outlined bars indicate methods that could not be evaluated because transcript-level coordinates were unavailable. Higher values indicate better performance for all metrics.

#### *A.3.3 cellAdmix*

NMF: number of factors was calculated as $\lceil 1.5 n_{\mathrm{CT}} \rceil$ where $n_{\mathrm{CT}}$ is the number of cell types represented in the dataset; $n_{\mathrm{CT}} = 15$ for the tested datasets. Factor annotation was done via "cell-bridging" as recommended by the authors [10].

#### *A.3.4 ovrlpy*

ovrlpy computes a vertical signal integrity (VSI) map, assigning integrity scores to pixels [11]. Corrected matrices for ovrlpy were computed via removal of transcripts at pixels with scores less than 0.7. This is identical to the method used in the SPLIT paper and follows the ovrlpy author recommendations in the following GitHub issue: https://github.com/HiDiHlabs/ovrl.py/issues/40

### A.4 Cell-type harmonization

#### *A.4.1 NSCLC CosMx Dataset*

To place the single-cell reference and spatial annotations in a shared ontology, we first integrated the reference with each spatial slice separately. For each slice, we concatenated the reference and that slice, normalized counts, applied $\log(1 + x)$ transformation and scaling, and computed 30 principal components. Harmony was then used to integrate the single-cell and spatial batches in PCA space [12].

Spatial labels were mapped directly to the shared cell-type ontology. For example, spatial labels for CD4 and CD8 memory or naive T cells were mapped to *T CD4+* and *T CD8+*, respectively; *mDC* and *pDC* were mapped to *DC*; and the annotated tumor subtypes were mapped to *Malignant*. Other labels, including macrophage, monocyte, fibroblast, endothelial, epithelial, mast, neutrophil, NK, B-cell, and plasmablast labels, were mapped to their corresponding shared categories.

Single-cell categories with a direct correspondence were also mapped directly. In particular, ciliated, club, type I, and type II cells were mapped to *Epithelial*; plasma cells to *Plasmablast*; neutrophils to *Neutrophil*; and patient-specific populations to *Malignant*. The remaining categories *T cells* and *MoMacDC* were subdivided using the spatial labels. For each spatial slice, a $k$-nearest-neighbor classifier with $k = 1$ and Euclidean distance was fitted in the slice-specific Harmony embedding. Single-cell *T cells* were assigned the label of their nearest spatial cell among *T CD4+*, *T CD8+*, and *Treg*; single-cell *MoMacDC* cells were assigned the label of their nearest spatial cell among *Macrophage*, *Monocyte*, and *DC*.

This procedure yielded one harmonized label for each reference cell in each slice-specific embedding. The final single-cell label was the modal label across slices:

$$\ell_i^{\text{consensus}} = \text{mode}_s\left(\ell_{is}\right),$$

where $\ell_{is}$ is the harmonized label assigned to cell $i$ using the embedding for slice $s$.

##### *A.4.2 Breast Cancer Xenium Dataset*

Label transfer from the scRNA-seq reference to Xenium was conducted using the same Harmony-integration + KNN strategy as described for the CosMx dataset.

#### A.5 Training details

SYNDRI models were trained independently for the whole-mouse-brain MERFISH, NSCLC CosMx, and breast-cancer Xenium datasets. Training examples were generated from pseudo-spatially mapped scRNA-seq profiles using the synthetic admixture procedure described in the main Methods. For the simulated NSCLC benchmark, a separate model was trained using the calibrated whole-gene donor-pool sampler described in Section A.8.

Each model was optimized for gene-level binary admixture classification using masked binary cross-entropy with logits. The positive class received a loss weight of 4.0. Optimization used AdamW with a configured learning rate of $2 \times 10^{-3}$, weight decay of 0.1, and gradient norm clipping at 1.0. The learning-rate schedule consisted of a linear warm-up followed by cosine decay. Training admixture examples were regenerated between epochs, whereas validation examples were held fixed.

Training used a data-loader batch size of 32 cells and gradient accumulation over five mini-batches, corresponding to an effective batch size of 160 cells per full accumulation window on one device. The models were trained in float32 precision with random seed 54. Full optimization and architecture hyperparameters are provided in Table S1, dataset-specific configurations and evaluated checkpoints in Table S2, and training and validation dataset sizes in Table S3.

Checkpoints were selected by inspecting validation AUPRC trajectories and identifying an early checkpoint after performance had reached a plateau. Automatic early stopping was not used. The evaluated checkpoints corresponded to milestones 130, 68, and 50 for the MERFISH, CosMx, and Xenium

**Table S1:** Training hyperparameters shared by the evaluated SYNDRI models. Dataset-specific settings and evaluated checkpoints are listed in Table S2.

| Parameter | Setting |
|---|---|
| Prediction task | Binary admixture classification per gene token |
| Transformer encoder layers | 4 |
| Embedding dimension | 128 |
| Attention heads | 8 |
| Feed-forward dimension | 512 |
| Activation | GELU |
| Hidden / attention dropout | 0.1 / 0.1 |
| Position embeddings | Learned BERT-style embeddings |
| Prediction head | Shared linear gene-token classification head |
| Optimizer | AdamW |
| Configured learning rate | $2 \times 10^{-3}$ |
| Adam coefficients | $\beta_1 = 0.9$, $\beta_2 = 0.95$ |
| Adam numerical stability | $\epsilon = 10^{-8}$ |
| Weight decay | 0.1 |
| Loss | Masked binary cross-entropy with logits |
| Positive-class loss weight | 4.0 |
| Data-loader batch size | 32 cells |
| Gradient accumulation | 5 mini-batches |
| Effective batch size | 160 cells for a full accumulation window on one device |
| Gradient norm clipping | 1.0 |
| Learning-rate schedule | Linear warm-up followed by cosine decay |
| Warm-up fraction | 0.1 of the configured scheduler horizon |
| Configured training horizon | 1,000 epochs |
| Numerical precision | Float32 |
| Random seed | 54 |
| Training order | Shuffled |
| Validation metrics tracked | Gene-level AUPRC, F1, AUROC |
| Zero-expression gene tokens | Dropped |
| Training admixture | Refreshed between epochs |
| Validation admixture | Fixed realization |
| Eligible admixture genes | Genes unexpressed in the native query |
| Same-type donors | Excluded |
| Minimum target addition | 10 transcripts |

models, respectively, and milestone 29 for the separate simulation-benchmark model (Table S2).

### A.6 Transcript localization analysis

To compare the spatial distribution of tumor-associated and T-cell-associated transcripts within annotated T cells, we analyzed 10,000 T cells sampled from the breast cancer Xenium dataset. Tumor-marker and T-cell-marker genes were defined using the accompanying annotation set; genes present in both categories were assigned to the T-cell-marker set.

Transcript localization was quantified relative to each cell boundary. For each marker transcript located within the corresponding cell polygon, we calculated the Euclidean distance to the cell boundary

**Table S2:** Dataset-specific SYNDRI training settings and checkpoints used for result generation. BM: whole-mouse-brain MERFISH; NC: NSCLC CosMx; BX: breast-cancer Xenium; SIM: the whole-gene donor-pool-fraction simulation benchmark. Optimizer updates are read directly from the saved checkpoint and agree with the optimizer's internal step counter. Milestones are zero-indexed epoch identifiers.

| Setting | BM | NC | BX | SIM |
|---|---|---|---|---|
| Evaluated checkpoint milestone | 130 | 68 | 50 | 29 |
| Epochs through evaluated checkpoint | 131 | 69 | 51 | 30 |
| Optimizer updates at checkpoint | 52,531 | 29,946 | 27,336 | 56,490 |
| Learning rate at checkpoint | $5.25 \times 10^{-4}$ | $2.76 \times 10^{-4}$ | $2.04 \times 10^{-4}$ | $1.20 \times 10^{-4}$ |
| Maximum input length (excluding `[CLS]`) | 1,536 | 1,536 | 512 | 1,536 |
| Configured gene context length | 1,530 | 1,530 | 506 | 1,526 |
| Donor neighbors | 5 | 5 | 5 | 9 |
| Minimum donor transcript count | 10 | 10 | 10 | 400 |
| Admixture generator | Whole-gene | Whole-gene | Whole-gene | Weighted whole-gene donor pool |
| Admixture parameter | $\tau \sim U(0.7, 0.85)$ | $\tau \sim U(0.7, 0.85)$ | $\tau \sim U(0.7, 0.85)$ | $\lambda \sim U(0, 0.25)$ |
| Calibrated donor-type weights | No | No | No | Yes |
| Input count clipping | None | None | Maximum 250 | None |
| Checkpoint used for evaluation | Explicit milestone | Explicit milestone | Explicit milestone | Fixed milestone after 30 epochs |

For BM, NC, and BX, the original sampler uses $\tau$ to specify a target final admixture fraction, with target addition $\mathrm{round}[\tau Q/(1-\tau)]$, subject to available eligible counts and whole-gene addition. For SIM, the target is $\mathrm{round}(\lambda P)$, where $P$ is the eligible, weighted donor-pool count. These parameters therefore have different meanings.
The 1,000-epoch setting defines the nominal scheduler horizon; it is not the number of epochs represented by the evaluated checkpoints. The scheduler computes $T = 1000\lfloor N_{\mathrm{train}}/32 \rfloor$ and uses $\lfloor 0.1T \rfloor$ warm-up steps. Its horizon does not divide by the gradient-accumulation factor. The evaluated checkpoints remain within this warm-up period. Validation F1 was tracked during training.

**Table S3:** Training and validation split sizes for the SYNDRI models.

| Dataset | Reference donors | Train cells | Validation cells |
|---|---|---|---|
| Whole mouse brain MERFISH | 222 | 64,013 | 15,976 |
| NSCLC CosMx | 7 | 69,382 | 18,224 |
| Breast Xenium | 4 | 85,616 | 22,453 |

and normalized this distance by the cell's equivalent radius,

$$r_{\mathrm{eq}} = \sqrt{\frac{A}{\pi}}, \qquad d_{\mathrm{norm}} = \frac{d_{\mathrm{boundary}}}{r_{\mathrm{eq}}},$$

where $A$ is the cell-polygon area. For each cell, normalized distances were averaged separately for tumor-marker and T-cell-marker transcripts. The resulting distributions therefore represent cell-level mean transcript localization rather than individual transcript distances.

We additionally quantified tumor-marker prevalence within each T cell as

$$q_i = \frac{N_{i,\text{tumor marker}}}{N_{i,\text{all transcripts}}},$$

where $N_{i,\text{tumor marker}}$ is the number of detected tumor-marker transcripts in cell $i$ and $N_{i,\text{all transcripts}}$ is its total detected transcript count. Cells with no tumor-marker transcripts were assigned $q_i = 0$. The distributions of $q_i$ and mean normalized transcript distance were used to compare tumor-marker abundance and intracellular localization with those of native T-cell markers.

#### A.7 Differential expression in tumor-associated and stromal fibroblasts

We evaluated how spillover correction affected differential expression between fibroblasts located in tumor and stromal regions of the CosMx NSCLC dataset. The analysis was adapted from the cellAdmix NSCLC full-data tutorial, with the modifications described below. We compared uncorrected expression (Raw) with expression corrected by SYNDRI, SPLIT, ResolVI, ovrlpy, and cellAdmix.

***Cell selection and matched analysis cohorts.*** Fibroblasts were identified using the harmonized cell-type annotation. The existing spatial niche annotations were retained: cells annotated as `tumor interior` were assigned to the tumor group, and cells annotated as `stroma` were assigned to the stromal group. Cells with other niche annotations were excluded from the differential expression analysis. Cell identities and region assignments were held fixed across correction methods.

Within each slice, we retained fibroblasts with at least 10 total counts and five detected genes in every available method, including Raw. The intersection of these passing cells defined a shared cohort for that slice. Consequently, the analysis compared expression on matched cells rather than allowing correction-dependent cell retention to change the comparison groups. Slice-level differential expression required at least 30 retained fibroblasts in each region. Corrected matrices were aligned to Raw by cell identifier and measured gene identity.

For the pooled differential expression analysis, we concatenated the retained cells from Lung5 replicates 1–3. Cells contributed equally, so replicates contributed in proportion to their retained cell counts. Raw, SYNDRI, SPLIT, ResolVI, and ovrlpy included 17,999 fibroblasts across all three replicates. Available cellAdmix results covered replicates 1 and 3, comprising 12,082 fibroblasts. For each Raw-versus-corrected $Z$-statistic comparison, Raw differential expression was recomputed using exactly the cells and slices available for that correction method.

***Corrected expression matrices.*** We used the previously generated corrected expression matrices for SPLIT, ResolVI, ovrlpy, and cellAdmix. The ovrlpy results used the correction output corresponding to the 0.7 VSI cutoff. ResolVI-derived expression estimates were analyzed numerically in the same downstream workflow as the other matrices; these estimates need not be integer transcript counts.

For SYNDRI, correction used hard zeroing of cell–gene entries based on predicted admixture probabilities. Thresholds were selected separately for each annotated cell type within each full slice, before selecting the fibroblast analysis cohort. Expressed entries with probability at least 0.5 were ranked by decreasing admixture probability. The threshold was chosen at the first cumulative count total reaching a target removal fraction of 0.5. All entries at or above that threshold were zeroed. If eligible entries contained less than the target number of counts, all eligible entries were removed. Whole-entry removal and tied probabilities could cause the achieved fraction to differ from the target. Counts with probabilities below 0.5 were protected.

***Differential expression testing.*** Each expression matrix was independently normalized to 10,000 total counts per cell and transformed using the natural logarithm:

$$Y_{ig}^{(m)} = \log\left(1 + 10^4 \frac{X_{ig}^{(m)}}{\sum_h X_{ih}^{(m)}}\right),$$

where $X_{ig}^{(m)}$ denotes expression of gene $g$ in cell $i$ under method $m$.

We compared tumor-region with stromal fibroblasts using the tie-corrected Wilcoxon rank-sum test implemented in `scanpy.tl.rank_genes_groups`. Testing was two-sided, and $p$-values were adjusted using the Benjamini–Hochberg procedure. All measured genes were retained as the multiple-testing background. Positive signed Wilcoxon statistics indicated higher expression in tumor-region fibroblasts; negative statistics indicated higher expression in stromal fibroblasts.

Volcano plots show Scanpy's approximate $\log_2$ fold change on the horizontal axis and $-\log_{10}$ of the BH-adjusted $p$-value on the vertical axis. The approximate fold change is calculated from back-transformed group means of log-normalized expression, rather than from a ratio of arithmetic mean raw counts. Reference lines indicate an absolute fold change of 1.5 and an adjusted $p$-value of 0.05. For display, fold changes were clipped to $[-5, 5]$, and adjusted $p$-values were floored at $10^{-30}$. These display limits did not alter the exported differential expression results.

***Marker-count removal.*** We summarized correction separately for malignant, macrophage, and native fibroblast marker sets. The malignant set comprised the 36-gene panel used in the cellAdmix tutorial. The macrophage set was *TYROBP, FCER1G, CD68, CSF1R, C1QA, C1QB, C1QC, CD163, MRC1, CD14*, and *ITGAM*. The native fibroblast set was *MYL9, CXCL12, PDGFRB, RARRES2*, and *CDH11*. Only measured genes contributed to each summary.

For marker set $M$ and method $m$, the fraction of Raw marker counts removed was

$$R_M^{(m)} = 1 - \frac{\sum_{i \in \mathcal{F}_m} \sum_{g \in M} X_{ig}^{(m)}}{\sum_{i \in \mathcal{F}_m} \sum_{g \in M} X_{ig}^{(\mathrm{Raw})}},$$

where $\mathcal{F}_m$ contains the matched fibroblasts from slices available for method $m$. This statistic uses pooled counts before library-size normalization. A value of zero indicates unchanged aggregate marker counts, one indicates complete removal, and negative values indicate increased estimated expression after correction.

The heatmap pooled all retained tumor- and stroma-region fibroblasts from eight slices: Lung5 replicates 1–3, Lung6, Lung9 replicates 1–2, Lung12, and Lung13. CellAdmix contributed seven available slices; its Raw denominator was restricted to those same slices.

***Changes in differential expression statistics.*** For each gene, we plotted the signed Wilcoxon statistic after correction against its corresponding Raw statistic. The diagonal $Z_{\mathrm{corrected}} = Z_{\mathrm{Raw}}$ denotes unchanged differential signal, whereas movement toward $Z_{\mathrm{corrected}} = 0$ indicates attenuation. Statistics were clipped to $[-20, 20]$ for visualization.

Malignant and macrophage markers were distinguished from other genes. The SYNDRI panels additionally annotated the tumor-associated fibroblast genes *IGFBP5, COL4A2, IGFBP3, FN1*, and *BGN*, and the stromal fibroblast genes *DCN, COL14A1, FOS, MMP2*, and *DUSP1.* These annotations did not affect testing or multiple-testing adjustment.

***Differences from the cellAdmix tutorial.*** Following the cellAdmix tutorial, we compared tumor-region and stromal fibroblasts using a Wilcoxon rank-sum framework. We used Scanpy's [13] tie-corrected implementation with explicit two-sided Benjamini–Hochberg adjustment, applying identical preprocessing and testing to all correction methods. Unlike the tutorial's Pagoda2-adjusted statistics, the Z comparisons show signed Wilcoxon test statistics before multiple-testing adjustment. This analysis constitutes a standardized adaptation of the tutorial rather than a numerical reproduction.

***Interpretation.*** The pooled tests treat individual cells as observations and do not model dependence among cells within a tissue section or spatial neighborhood. The Lung5 replicates originate from one donor. Accordingly, the adjusted $p$-values and changes in $Z$ statistics describe within-dataset differential expression and its response to correction; they do not establish differential expression across independent biological donors. The differential-expression analysis is adapted from the cellAdmix paper [10].

### A.8 SYNDRI-SIM

We constructed SYNDRI-SIM to evaluate spillover correction against known native and added expression counts. Single-cell RNA-seq expression profiles were assigned to observed CosMx cell positions, after which synthetic spillover was introduced from neighboring cells. This construction separates the expression used to generate the benchmark from the real spatial measurements used to assess distributional similarity.

***Reference partitions and pseudo-spatial mapping.*** The NSCLC single-cell reference was partitioned by patient: patient p5 supplied validation cells, patient p1 supplied test cells, and the remaining patients supplied training cells. Training cells were mapped independently to Lung5_Rep1–3, validation cells to Lung6, and test cells to Lung9_Rep1, Lung9_Rep2, Lung12, and Lung13. Regulatory T cells were excluded from both the reference and spatial data.

For each partition–slice pair, reference and spatial profiles were integrated using Harmony with 30 principal components. Reference cells were then assigned to unique spatial recipient positions using the integrated representation, requiring matching harmonized cell-type annotations. Candidate matching used up to 50 reference neighbors, a reuse penalty of 9, and random seed 0. Reference expression profiles could be reused across recipient positions, but reference cells were not shared across training, validation, and test partitions. The resulting pseudo-spatial data retained single-cell expression counts and the corresponding spatial recipient coordinates and annotations. Analyses used the common panel of 956 genes.

***Spatial donor neighborhoods.*** For each pseudo-spatial query cell $i$, we selected up to nine nearest eligible donor cells from the same slice, excluding donors with the same annotated cell type as the query. Eligible donors contained at least 400 native transcripts. Neighborhoods were constructed from the mapped spatial coordinates and held fixed across simulation candidates.

Let $X_{ig}$ denote the native count of gene $g$ in query $i$, $t_i$ its cell type, and $\mathcal{N}(i)$ its selected donor neighborhood. Synthetic additions were restricted to genes absent from the native query, $X_{ig} = 0$. Donor expression always referred to the original native counts; previously added contamination was not propagated between cells.

***Calibration of donor-type weights.*** We estimated a global contribution weight for each donor cell type using only Lung5_Rep1–3. Marker sets were obtained from cell-type-versus-rest DESeq2 results, retaining genes with adjusted $p \leq 0.05$, positive log fold change, and significant positive enrichment for exactly one cell type. This produced 267 markers across 14 cell types.

For donor type $d$, let $M_d$ denote its marker set and

$$n_{i,d} = |\{j \in \boldsymbol{\mathcal{N}}(i) : t_j = d\}| \,.$$

Queries were considered exposed to donor type $d$ when $n_{i,d} > 0$. Donor-marker availability was calculated as

$$E_{i,d} = \frac{\sum_{\substack{j \in \boldsymbol{\mathcal{N}}(i) \\ t_j = d}} \sum_{g \in M_d} X_{jg} \, \mathbf{1}\{X_{ig} = 0\}}{|M_d| \, n_{i,d}} \,.$$

Thus, availability measured eligible native marker counts per marker and per selected donor, conditional on donor presence.

For each calibration slice $s$ and recipient type $r \neq d$, we measured the average detection frequency of donor markers in the matched real CosMx recipients exposed to $d$:

$$C^{\mathrm{ST}}_{s,r,d} = \frac{1}{|\boldsymbol{\mathcal{I}}_{s,r,d}| \, |M_d|} \sum_{i \in \boldsymbol{\mathcal{I}}_{s,r,d}} \sum_{g \in M_d} \mathbf{1}\{Y^{\mathrm{real}}_{ig} > 0\},$$

where $\boldsymbol{\mathcal{I}}_{s,r,d}$ contains the corresponding mapped recipients with $n_{i,d} > 0$. Exposure was defined using the same selected donor graph as the pseudo-spatial data. We compared this quantity with marker detection in reference cells of recipient type $r$, denoted $C^{\mathrm{ref}}_{r,d}$, and calculated

$$\Delta_{s,r,d} = \max\bigl(0, \, C^{\mathrm{ST}}_{s,r,d} - C^{\mathrm{ref}}_{r,d}\bigr) \,.$$

Excess detection and marker availability were averaged equally across supported recipient types within each slice, then equally across calibration slices. Strata required at least 50 donor-exposed mapped and matched spatial recipients and at least 50 reference cells of the recipient type. Donor types required support from at least two calibration slices and positive marker availability.

For supported donor types, the unnormalized weight was

$$u_d = \frac{\overline{\Delta}_d}{\overline{E}_d + 10^{-8}} .$$

These weights were normalized to mean one and shrunk 25% toward one:

$$w_d = 0.75 \frac{u_d}{\mathrm{mean}_{d' \in \boldsymbol{\mathcal{D}}_{\mathrm{sup}}} u_{d'}} + 0.25.$$

Unsupported donor types received the neutral weight $w_d = 1$. Weights were frozen before test-set simulation. They represent effective contributions inferred from cross-platform marker detection, rather than directly measured transcript-transfer probabilities. Weights were not renormalized within individual neighborhoods.

***Whole-gene donor-pool sampling.*** Weighted donor contributions were converted to integer counts by stochastic rounding:

$$Z_{j\to i,g} = \lfloor w_{t_j} X_{jg} \rfloor + B_{ijg}, \qquad B_{ijg} \sim \mathrm{Bernoulli}\left(w_{t_j} X_{jg} - \lfloor w_{t_j} X_{jg} \rfloor\right).$$

The eligible pooled count for gene $g$ was

$$P_{ig} = \mathbf{1}\{X_{ig} = 0\} \sum_{j \in \boldsymbol{\mathcal{N}}(i)} Z_{j\to i,g}, \qquad P_i = \sum_g P_{ig}.$$

For each query, a transfer fraction was sampled as

$$\lambda_i \sim U(0, \lambda_{\max}), \qquad T_i = \mathrm{round}(\lambda_i P_i).$$

Queries with an empty eligible pool or a target below 10 transcripts received no addition. Otherwise, genes with $P_{ig} > 0$ were randomly permuted, and each selected gene's entire pooled count was added until the cumulative addition reached or exceeded $T_i$. The final gene was not truncated. Consequently, the achieved addition can exceed the target.

Writing $A_{ig}$ for the added counts, the simulated expression was

$$Y_{ig} = X_{ig} + A_{ig}.$$

The realized contamination fraction was recorded as

$$f_i = \frac{\sum_g A_{ig}}{\sum_g X_{ig} + \sum_g A_{ig}}.$$

Here, $\lambda_i$ controls the fraction of the eligible weighted donor pool targeted for transfer; it is not the final contamination fraction $f_i$. Donation was additive and did not deplete source cells. Native, added, combined, and donor–recipient–gene allocations were retained separately.

***Selection of transfer efficiency.*** We evaluated

$$\lambda_{\max} \in \{0.1, 0.25, 0.5, 0.75, 1.0\}$$

using simulation seeds 0, 1, and 2. A no-admixture baseline was evaluated separately. Parameter selection used only the three Lung5 calibration slices, with up to 500 fixed queries per supported cell type and slice. Supported types required at least 50 mapped recipients and 50 reference cells. The same queries were used across candidates and seeds.

Expression vectors were normalized to 10,000 counts per cell and transformed by $\log(1+x)$. Within each slice and cell type, we compared simulated and matched real expression distributions using the biased squared maximum mean discrepancy:

$$\widehat{\mathrm{MMD}}^2(U, V) = \frac{1}{n^2} \sum_{i,j=1}^{n} k(u_i, u_j) + \frac{1}{m^2} \sum_{i,j=1}^{m} k(v_i, v_j) - \frac{2}{nm} \sum_{i=1}^{n} \sum_{j=1}^{m} k(u_i, v_j),$$

with

$$k(u, v) = \exp\left(-\frac{\|u - v\|_2^2}{2\sigma^2}\right).$$

The bandwidth $\sigma^2$ was the median positive squared pairwise distance among pooled clean pseudo-spatial and matched real expression vectors. It was fixed within each slice–type stratum across all candidates and seeds.

Losses were averaged equally across supported cell types within each slice, then across slices and simulation seeds. This selected $\lambda_{\max} = 0.25$. Detected-gene counts, non-native marker detection, and cross-type marker co-detection were reported as separate diagnostics and were not combined with $\text{MMD}^2$ to select the parameter. Held-out test slices were not used for weight estimation or parameter selection.

For Fig. 3a, each boxplot observation is a cell-type-specific $\text{MMD}^2$ value from one calibration slice. After-admixture values are averaged across the three simulation seeds. These statistics compare distributions of individual cell expression vectors, rather than cell-type pseudobulks.

***Cell-type adjacency.*** For the selected test slice, we constructed directed six-nearest-neighbor graphs from cell centroids. Let $N_{rd}$ be the number of directed edges from query type $r$ to neighbor type $d$, $N$ the number of cells, and $\pi_r$ the overall abundance of type $r$. Adjacency enrichment was

$$H_{rd} = \log_2\left[\frac{N_{rd}/(6N) + 10^{-8}}{\pi_r \pi_d + 10^{-8}}\right].$$

With a fixed neighborhood size, this is the abundance-adjusted enrichment of neighbor type $d$ around query type $r$.

For each ordered pair of type indices $r < d$, the main triangular panel places simulated $H_{rd}$ above the diagonal and the same directed real statistic $H_{rd}$ at the mirrored lower position. Reverse-direction values are displayed separately in the notebook; diagonal entries are the mean of real and simulated values. Because the construction retains the matched recipient positions and labels, agreement in adjacency is expected by design.

***Marker-gene co-detection.*** For Fig. 3c, we selected three shared marker genes per cell type from the DESeq2 candidate-marker table, ranked by increasing adjusted $p$ value and then decreasing log fold change. This display panel was distinct from the full exclusive-marker set used for donor-weight calibration.

For each gene pair $(g, h)$, co-detection was

$$Q_{gh} = \frac{1}{N}\sum_{i=1}^{N} \mathbf{1}\{X_{ig} > 0\}\mathbf{1}\{X_{ih} > 0\}.$$

Every matrix entry used all cells in the corresponding dataset, irrespective of their annotated type. We compared the unique training/validation single-cell reference, the simulated test slice, and its exactly matched real spatial recipients. The displayed configuration used raw joint-detection fractions, without row normalization or logarithmic transformation.

***Correction methods.*** All methods were evaluated on the same saved simulated test matrices. SYNDRI was retrained using the original training/validation partitions and training hyperparameters, with the whole-gene $\lambda_{\max} = 0.25$ sampler replacing the previous admixture generator. Inference used checkpoint milestone 29.

ResolVI was fitted independently to each test slice for 100 epochs; corrected expression was obtained from the posterior median of `px_rate` using three posterior samples. RCTD annotations and SPLIT correction used the unique training/validation single-cell reference, excluding duplicate reference profiles introduced by pseudo-spatial remapping. No test-reference cells were included in this reference.

For transcript-based methods, simulated native transcripts were placed uniformly within each recipient's segmentation mask, with random subpixel offsets. Added transcripts were also placed inside the recipient mask, with pixel-sampling weights proportional to

$$\exp[-d(x, P_j)/(2\,\mu\mathrm{m})]\,,$$

where $P_j$ is the known donor segmentation region. Discrete z-planes were sampled from the empirical slice-wide distribution. Transcript counts and donor identities followed the saved simulation allocations exactly.

The cellAdmix workflow inferred transcript factors, annotated type-specific factors using its bridge and false-positive checks, and reconstructed corrected counts after removing the selected assignments. Ovrlpy was fitted independently per FOV; its evaluated corrected counts removed transcripts with pixel-level VSI below 0.7. Transcript ground-truth labels were excluded from method fitting.

***Precision–recall benchmarking.*** Evaluation considered only observed cell–gene entries,

$$\boldsymbol{\mathcal{E}} = \{(i, g) : Y_{ig} > 0\},$$

with ground-truth label

$$y_{ig} = \mathbf{1}\{A_{ig} > 0\}.$$

Because additions were restricted to native-zero genes, positive entries contained exclusively added expression.

For SYNDRI, the ranking score was the predicted admixture probability $p_{ig}$. For methods supplying corrected counts $C_{ig}$, the score was

$$s_{ig} = 1 - \frac{C_{ig}}{Y_{ig}}.$$

Thus, larger scores indicate greater removal of the observed expression. Ovrlpy's gene-level score was derived from its reconstructed corrected counts, rather than from a broadcast cell-level integrity score.

Labels and scores were pooled across all four held-out test slices before calculating precision–recall curves. Each observed cell–gene entry contributed equally; entries were not weighted by transcript abundance. AUPRC was computed by trapezoidal integration, with average precision recorded separately. The no-skill precision baseline was the positive-entry prevalence,

$$\pi = \frac{\sum_{(i,g)\in\boldsymbol{\mathcal{E}}} y_{ig}}{|\boldsymbol{\mathcal{E}}|}.$$

For each method, the annotated operating point maximized

$$F_1(t) = \frac{2\ \mathrm{Precision}(t)\,\mathrm{Recall}(t)}{\mathrm{Precision}(t) + \mathrm{Recall}(t)}.$$

Binarized variants used the corresponding threshold $t^\star$ to remove all counts when $s_{ig} \geq t^\star$ and retain all counts otherwise. Their precision–recall curves therefore contain only one nontrivial operating point, together with the standard endpoints.

Best-$F_1$ thresholds were selected on the pooled test labels for descriptive comparison; they are not independently validated deployment thresholds. Metrics were calculated from full-resolution scores. Curve coordinates were subsequently reduced for plotting, retaining the best-$F_1$ point.

### A.9 Context response experiment

We evaluated whether SYNDRI's gene-level admixture predictions respond to neighborhood composition when the query cell's expression and annotated identity are held fixed. The experiment used eight breast-cancer Xenium sections (B1_1, B1_2, B1_3, B5, B9, B10, B12, and B17) and a fixed, pretrained breast-cancer model. No model parameters were updated during this experiment.

***Query cells and context perturbations.*** For each cell, the original context consisted of the cell-type tokens of its five nearest eligible neighbors with a different annotated cell type. Eligible query cells had finite spatial coordinates, a valid cell-type annotation, at least 10 measured transcripts, at least five detected genes, and a complete five-neighbor context.

Within each section and exact query cell type, we selected up to 300 query cells using the stratified design described below. Alternative contexts were obtained from other eligible cells of the same exact type in the same section. We selected up to five distinct context-source cells for each possible malignant-neighbor count, $m \in \{0, \ldots, 5\}$, giving at most 30 alternative contexts per query type and section. Sampling was without replacement; unavailable context-count categories were not filled using other categories. Distinct source cells were retained even when they supplied identical token sequences.

Each query was evaluated under the selected alternative contexts while retaining its original expression vector and query-type token. Only the neighborhood cell-type tokens were replaced; expression profiles from the context-source cells were not transferred. We also evaluated the original context and an unchanged-context duplicate as a consistency control. Unchanged duplicates were excluded from response summaries.

***Query cell type families.*** Exact query annotations were grouped into four cell type families for visualization:

- *Lymphoid*: CD4 memory T cells, cytotoxic T cells, regulatory T cells, natural killer cells, and plasma cells;
- *Myeloid*: classical monocytes, macrophages, mast cells, and plasmacytoid dendritic cells;
- *Stromal*: breast fibroblasts, lung fibroblasts, and smooth-muscle cells;
- *Vascular*: capillary endothelial cells.

The model continued to receive the exact cell-type annotation. Cell type families were used only to summarize responses. Pericytes were excluded from the matched-native-marker analysis because the published marker annotation did not provide a pericyte marker set; no proxy marker set was substituted.

***Malignant and native marker sets.*** Let $M$ denote the fixed 32-gene breast-cancer panel, and let $N_t$ denote the published native-marker set assigned to exact query type $t$, using Supplementary Table 4 of the source Xenium study. The three T-cell annotations shared the published T-cell marker set, the two fibroblast annotations shared the general fibroblast set, and plasmacytoid dendritic cells used the general dendritic-cell set. Plasma cells used the explicitly annotated plasma-cell marker, *TNFRSF17*.

For query $i$ with observed expression $X_i$, we restricted comparisons to measured, expressed genes:

$$M_i = \{g \in M : X_{ig} > 0\}, \qquad N_i = \{h \in N_{t_i} : X_{ih} > 0\}.$$

These sets were fixed across all context perturbations of that query. Queries without at least one expressed malignant-panel gene and one expressed native marker were excluded from score-based analyses. The malignant and native sets were disjoint.

***Marker removal priority.*** Let $p_{ig}(c)$ be the predicted admixture probability for gene $g$ in query $i$ under supplied context $c$. We defined malignant-marker removal priority as the AUROC comparing malignant-panel genes with native markers:

$$A_i(c) = \frac{1}{|M_i|\,|N_i|} \sum_{g \in M_i} \sum_{h \in N_i} \left[ \mathbf{1}\{p_{ig}(c) > p_{ih}(c)\} + \frac{1}{2}\mathbf{1}\{p_{ig}(c) = p_{ih}(c)\} \right].$$

Thus, $A_i(c)$ is the probability that a randomly selected expressed malignant-panel gene receives a higher admixture score than a randomly selected expressed native marker, with half credit for ties. Larger values indicate preferential ranking of malignant-panel genes for removal relative to native expression. This is a ranking statistic, rather than a fraction of transcripts removed or an accuracy measurement against known contamination labels.

Writing $c_i^0$ for the original context, we calculated

$$\Delta A_i(c) = A_i(c) - A_i(c_i^0), \qquad \Delta m_i(c) = m(c) - m(c_i^0),$$

where $m(c)$ is the number of malignant-cell tokens among the five supplied neighbors. Positive $\Delta A_i$ indicates increased relative removal priority under the substituted context. Context swaps with $\Delta m_i = 0$ can still change predictions because the identities of the nonmalignant neighbors may differ.

***Nine query strata.*** To distinguish context response from baseline expression and neighborhood composition, queries were stratified along two axes. First, malignant-marker burden was defined from the original counts as

$$B_i = \frac{\sum_{g \in M} X_{ig}}{\sum_h X_{ih}}.$$

Eligible cells were divided into low, medium, and high burden groups using tertiles calculated separately within each section and exact cell type. Second, original malignant-neighbor counts were grouped as $0$, 1–2, or 3–5. Their Cartesian product defined nine possible query strata.

The query-sampling quota was distributed approximately equally across available strata, subject to their capacities. Burden cutoffs were not recalculated after filtering for expressed native markers. Consequently, some strata could have insufficient support or be absent.

***Cell type family response curves.*** For Fig. 5b, responses from context-source cells giving the same $\Delta m_i$ were first averaged within each query. We then calculated the median query response within each section, exact type, burden group, original-context stratum, and $\Delta m$ value, requiring at least five contributing queries. Supported exact-type summaries were averaged equally within each cell type family.

Original-context strata were retained only where supported by at least three sections, then averaged equally within each section at each $\Delta m$. Curves show the median of these section-level summaries; shaded regions show their interquartile range. Sections and supported original-context strata therefore contribute equally rather than in proportion to their numbers of cells.

***Gene-specific context response and missing heatmap entries.*** For each expressed malignant-panel gene, we additionally defined its removal priority relative to native markers as

$$W_{ig}(c) = \frac{1}{|N_i|} \sum_{h \in N_i} \mathbf{1}\{p_{ig}(c) > p_{ih}(c)\}.$$

Unlike the panel-level AUROC, this gene-specific statistic uses strict comparisons, so ties contribute zero. We calculated

$$\Delta W_{ig}(c) = W_{ig}(c) - W_{ig}(c_i^0)$$

and, separately within each section, cell type family, and query stratum,

$$\rho_{s,F,b,o,g} = \mathrm{Spearman}\left(\Delta m_i(c),\, \Delta W_{ig}(c)\right).$$

A correlation required at least 10 distinct eligible queries, at least three distinct $\Delta m$ values, and a nonconstant response. Fig. 5c displays the median correlation across supported section–stratum combinations, requiring support from at least three sections.

Positive correlations indicate that increasing malignant context raises a gene's removal priority relative to native markers; negative correlations indicate the reverse. Gray or blank entries represent insufficient support or undefined correlations, including constant responses. They do not represent zero response or nonsignificant effects. White entries near the color-scale midpoint indicate supported correlations near zero.

***Controlled and original-context response slopes.*** We compared the response induced by context substitution with the association observed under original spatial contexts (Fig. 5d). For each fixed query, we fitted an ordinary least-squares regression with an intercept:

$$\Delta A_i(c) = \alpha_i + \beta_{\mathrm{ctrl},i} \Delta m_i(c) + \varepsilon_{ic}.$$

Queries required at least three distinct context-count levels. The coefficient $\beta_{\mathrm{ctrl},i}$ measures the change in malignant-marker removal priority per additional malignant-neighbor token while query expression and identity remain fixed.

For the same eligible queries, we fitted a weighted regression within each section, exact query type, and burden group:

$$A_i(c_i^0) = a_{s,t,b} + \beta_{\mathrm{native},s,t,b}\, m(c_i^0) + \epsilon_i.$$

Here, $\beta_{\mathrm{native}}$ measures the association between original malignant-neighbor count and removal priority across different cells of the same exact type. "Native" refers to the original, unperturbed context, not to known uncontaminated expression.

Original-context strata contributed equal total weight within each fit, with equal weights for queries within a stratum. Type–stratum combinations required at least five eligible queries; original-context regressions required at least 10 queries and three distinct malignant-neighbor counts. Controlled slopes were averaged using the same query weights. Only exact-type groups supplying both valid coefficients contributed to cell type family comparisons.

Exact-type coefficients were averaged equally within each section and cell type family. Small points in Fig. 5d represent section-level coefficient pairs; large points and error bars show coordinate-wise medians and interquartile ranges across sections.

Positive $\beta_{\text{ctrl}}$ indicates that supplying more malignant neighbors increases relative malignant-marker removal priority. Positive $\beta_{\text{native}}$ indicates a corresponding association in unperturbed tissue. Agreement in sign supports consistency between the controlled model response and the observed spatial association. However, $\beta_{\text{native}}$ remains observational and can reflect differences between cells beyond their neighborhood composition. Neither coefficient establishes that the affected transcripts are truly contaminating molecules. The plotted interquartile ranges describe section-to-section variability and are not confidence intervals or significance tests.

***Vascular expression plausibility.*** To examine the heterogeneous vascular response (Fig. 5e), we fitted a context-response slope for each expressed panel gene in each eligible capillary endothelial query:

$$W_{ig}(c) = a_{ig} + \beta_{ig}\Delta m_i(c) + e_{igc}.$$

Queries again required at least three context-count levels. Each gene's vascular response was summarized by the median slope across eligible queries.

Using the matched single-cell reference, we measured expression prevalence in capillary endothelial and malignant cells. With $k_{g,t}$ expressing cells among $n_t$ reference cells, the plotted plausibility contrast was

$$L_g = \log_2\left[\frac{(k_{g,\text{vascular}} + 0.5)/(n_{\text{vascular}} + 1)}{(k_{g,\text{malignant}} + 0.5)/(n_{\text{malignant}} + 1)}\right].$$

Positive values indicate greater detection prevalence in vascular cells; negative values indicate greater prevalence in malignant cells. Comparing $L_g$ with the vascular context-response slope assesses whether genes with plausible endogenous vascular expression respond differently from genes preferentially detected in malignant cells.

The association between gene-specific vascular context-response slopes and the smoothed $\log_2$ detection-prevalence ratio was quantified using Spearman rank correlation across 32 genes. Statistical significance was assessed using a two-sided permutation test with 10,000 permutations, randomly permuting the response slopes across genes. The permutation $p$-value was calculated as

$$p_{\text{perm}} = \frac{1 + \sum_{b=1}^{10{,}000} \mathbf{1}(|\rho_b| \geq |\rho_{\text{obs}}|)}{10{,}001}.$$

### A.10 Scalable execution of cellAdmix

To make cellAdmix tractable on the transcript-level SYNDRI-SIM datasets, we modified its execution and data-storage strategy while retaining its NMF, conditional random field (CRF), and factor-annotation procedures. The adaptations comprised cell-preserving CRF batching, streamed construction of annotation inputs, incremental reduction of transcript neighborhoods, and reuse of factor-independent geometric calculations. The installed cellAdmix package was retained unchanged, and the original execution path remained available.

***Shared factorization and cell-preserving CRF batching.*** We fitted one NMF model per slice, using $K = \lceil 1.5C \rceil$ factors for $C$ annotated cell types ($K = 21$ for 14 types). The existing cell-type-balanced sampling procedure used a target of 1,000 cells, a transcript-neighborhood size of 20, the `nmol.dsamp=2000` setting, and random seed 0. The fitted model was checkpointed and shared across all subsequent CRF batches; separate models were not fitted to individual batches.

Because cellAdmix performs CRF inference separately within each cell, we partitioned inference into batches containing at most 128 complete cells and 250,000 transcripts. A cell exceeding the transcript limit was processed intact, preserving its within-cell graph. Each batch called the original `run_crf_all` implementation with 10 transcript neighbors in a fresh R process, allowing temporary worker memory to be released after completion. The parent process retained the input table and a single factor-assignment vector rather than retaining all intermediate CRF objects.

Before batching, we determined the complete gene universe among CRF-eligible cells and applied the original missing-gene loading augmentation once to the shared NMF model. Freezing this augmentation prevented batch-specific gene coverage from changing loading normalization. The CRF graph construction, potentials, transition settings, and iteration limits were otherwise inherited from cellAdmix. Assignments were restored to the original molecule order and checked for completeness.

***Streamed annotation inputs.*** Transcript records and their CRF assignments were read together in chunks of 250,000 rows. Molecule identifiers, cell identifiers, genes, and cell-type annotations were checked for agreement between the two inputs. Coordinates and assignments were stored in memory-mapped binary arrays, ordered by cell while preserving the original within-cell molecule order. Cell–factor counts and transcript-derived cell centroids were accumulated during streaming. Transcripts lacking CRF assignments were excluded consistently with the annotation input, and their number was recorded explicitly.

***Incremental transcript-neighborhood reduction.*** The original annotation workflow materializes transcript-level neighborhood information before reducing it to candidate cell pairs. We instead constructed one exact three-dimensional `cKDTree` over the assigned transcripts and queried it in chunks of 100,000 transcripts. Each query retained its 10 nearest other transcripts, explicitly excluding the query transcript itself. All fields of view remained in the same spatial index, preserving potential interactions across field-of-view boundaries.

Neighborhood results were immediately reduced to the cell-pair information consumed by the original annotation procedure. The reduction preserved its first-qualifying-molecule, lexical adjacent-cell selection, same-type eligibility, and duplicate-removal rules. The complete transcript-neighbor graph was therefore unnecessary. No additional transcript, cell, or field-of-view downsampling was introduced at this stage.

***Bounded factor annotation and geometry reuse.*** Bridge tests were evaluated serially in the original cell-type-pair order using random seed 0. Cell–factor fractions were obtained from the accumulated count summaries. We applied the original cell-pair pruning procedure with a 200-cell sampling setting before loading transcript coordinates, so only the geometry needed for the selected cells was materialized for each test.

For a fixed cell pair and coordinate configuration, nearest-neighbor geometry is shared across factor-specific bridge scores. We therefore cached the original RANN-based neighborhood calculations and factor-specific boundary-crossing fractions, reusing them across factors. These deterministic calculations consume no random numbers. The original null-generation procedure, including its geometric transformations and random draws, was retained.

We preserved three null iterations, a bridge-neighborhood size of 20, the one-sided paired Wilcoxon test, and the 75th-percentile observed bridge statistic. Candidate factors were extracted using the original joint-NMF procedure with an unadjusted significance threshold of 0.1. The distant-cell false-positive check used the accumulated cell–factor fractions and a cell-centroid graph with up to 100 neighbors, retaining the original median threshold of 0.1. Synthetic contamination labels were not used to select factors.

***Checkpointing and reproducibility.*** NMF fits, completed CRF batches, neighborhood-reduction progress, and individual bridge tests were checkpointed. Bridge checkpoints included the random-number-generator state, allowing completed tests to be reused while preserving the subsequent random stream. Input signatures and configuration checks guarded checkpoint reuse; the annotation cache additionally recorded implementation hashes.

These changes bound temporary neighborhood-query and CRF-worker allocations. They do not make total memory independent of dataset size: the coordinate index, staging-order arrays, and parent CRF input still require storage that scales with the number of transcripts.

***Validation against the original implementation.*** We compared the annotation adaptation with the original single-worker cellAdmix implementation using seed 0 on a constructed dataset containing 120 cells and 8,400 transcripts and a simulation-derived Lung12 subset containing 60 cells and 67,118 transcripts. Global neighbor sets, selected cell pairs, cell–factor counts, bridge statistics, candidate annotations, false-positive filtering, and final factor maps agreed on both datasets; centroids agreed within $10^{-12}$. Because both subset comparisons produced empty final factor maps, an additional constructed case verified agreement for a retained factor–cell-type assignment and rejection of a native factor.

These checks establish agreement on the evaluated subsets, rather than universal bitwise equivalence on complete slices. In particular, exact-distance ties may be ordered differently by the SciPy [14] and FNN nearest-neighbor implementations.
